# Neurochemistry-enriched dynamic causal models of magnetoencephalography, using magnetic resonance spectroscopy

**DOI:** 10.1101/2022.06.17.493881

**Authors:** Amirhossein Jafarian, Laura E Hughes, Natalie E Adams, Juliette Lanskey, Michelle Naessens, Matthew A Rouse, Alexander G Murley, Karl J Friston, James B Rowe

## Abstract

We present a hierarchical and empirical Bayesian framework for testing hypotheses about synaptic neurotransmission, based on the integration of ultra-high field magnetic resonance spectroscopy (7T-MRS) and magnetoencephalography data (MEG). A first level dynamic causal modelling of cortical microcircuits is used to infer the connectivity parameters of a generative model of individuals’ neurophysiological observations. At the second level, individuals’ 7T-MRS estimates of regional neurotransmitter concentration supply empirical priors on synaptic connectivity. We compare the group-wise evidence for alternative empirical priors, defined by monotonic functions of spectroscopic estimates, on subsets of synaptic connections. For efficiency and reproducibility, we used Bayesian model reduction (BMR), parametric empirical Bayes and variational Bayesian inversion. In particular, we used Bayesian model reduction to compare models of how spectroscopic neurotransmitter measures inform estimates of synaptic connectivity. This identifies the subset of synaptic connections that are influenced by neurotransmitter levels, as measured by 7T-MRS. We demonstrate the method using resting-state MEG (i.e., task-free recording) and 7T-MRS data from healthy adults. We perform cross-validation using split-sampling of the MEG dataset. Our results confirm the hypotheses that GABA concentration influences local recurrent inhibitory intrinsic connectivity in deep and superficial cortical layers, while glutamate influences the excitatory connections between superficial and deep layers and connections from superficial to inhibitory interneurons. The method is suitable for applications with magnetoencephalography or electroencephalography, and is well-suited to reveal the mechanisms of neurological and psychiatric disorders, including responses to psychopharmacological interventions.

## 1 Introduction

This paper introduces an empirical Bayesian methodology for inferring synaptic physiology from human, *in vivo* neurophysiological recordings and *in vivo* measurements of neurochemicals, including neurotransmitters. It is an example of a broader class of enriched dynamic causal models (DCMs), informed by multimodal data that can incorporate neurochemistry, molecular pathology or selective loss of neuronal sub-populations and their synapses. Magnetic resonance spectroscopy (MRS) can be used to estimate individual differences in key neurochemical concentrations in local brain regions (Blüml and Panigrahy, 2012). It has been used to quantify neurochemical changes in neurological and neuropsychiatric disorders, such as Alzheimer’s disease (Wang et al., 2015), frontotemporal lobar degeneration-related syndromes (Murley et al., 2020, Murley et al., 2022), and schizophrenia (Jelen et al., 2018, Cohen et al., 2015, Duarte and Xin, 2019), as well as normal aging (Boumezbeur et al., 2010). Such spectroscopy can be predictive of task performance and the response to pharmacological interventions (Chowdhury et al., 2012, Stagg et al., 2011, McColgan et al., 2020, Gawne et al., 2020, Schmitz et al., 2017, Adams et al., 2021). Differences in neurotransmitter levels influence synaptic transmission, which in turn affects the generators of magneto- and electro-encephalographic signals (M/EEG).

Here, we characterise the relationship between physiology and neurotransmitter levels, using a hierarchical Bayesian approach (Friston et al., 2016). We present a method for specifying and comparing the evidence for hierarchical models (using parametric empirical Bayes) that encodes the effect of neurotransmitter levels (as measured with MRS) on different combinations of synaptic parameters in a neural mass model of MEG recordings. Our approach complements previous studies, where correlational analysis has been used to investigate relationships between spectroscopy measures and electrophysiology (Rideaux, 2021, Steel et al., 2020, Kober et al., 2001). We envisage that our method may be useful for characterising the synaptic deficits in several neuropsychiatric conditions (e.g., (Adams et al., 2021, Limongi et al., 2021)).

MRS is a non-invasive neuroimaging modality used to estimate biochemical concentrations, including the neurotransmitters glutamate and gamma-aminobutyric acid (GABA) (McColgan et al., 2020, Stagg and Rothman, 2013). Glutamatergic neurotransmission mechanisms include release, reuptake into astrocytes, conversion to glutamine then glutamate, and vesicular repacking (Gruetter et al., 2001, Pellerin et al., 2007, van der Graaf, 2010). In contrast, GABA cycling is predominantly neuronal where, following a release phase, there is presynaptic GABA reuptake and vesicular repackaging (Gruetter et al., 2001). The majority of GABA and glutamate is intracellular (Myers et al., 2014), but total GABA and glutamate concentrations correlate with neurophysiological features such as gamma oscillatory power and corticospinal excitability (Lally et al., 2014, Greenhouse et al., 2017).

In what follows, we show that neurotransmitter levels can be used as prior constraints on the estimation of effective synaptic connectivity, in biophysically-informed models of cortical function. For example, glutamate and GABA concentrations control the dynamics of the excitatory amino-3-hydroxy-5-methyl-4-isoxazolepropionic acid (AMPA) receptors and inhibitory GABAergic receptors respectively (Rideaux, 2020, Gruetter et al., 2001). The aim of the current work was to compare the evidence for alternate hypotheses regarding the relationships between glutamate/GABA concentrations and synaptic connectivity, quantified using non-invasive MRS recordings and M/EEG data, respectively.

The first step in establishing this relationship is to infer synaptic parameters from neurophysiological observations. Since Hodgkin and Huxley (Hodgkin and Huxley, 1952), there have been many approaches to examine micro-, meso- and macroscale brain dynamics with many options for model identification (Nelson and Rinzel, 1998, Steyn-Ross and Steyn-Ross, 2010, Robinson et al., 2003, Terry et al., 2022, Jirsa et al., 2014, Deco et al., 2008). DCMs build directly on the modelling framework established by Hodgkin and Huxley, who proposed a linear dynamical system model to explain the relation between conductance dynamics and ion current dynamics. Later inclusion of nonlinear dynamics improved performance, balancing complexity with accuracy in a way that remains highly relevant to DCM (e.g., (Nelson and Rinzel, 1998)). The hypothesis testing machinery in DCM—enabling inferences on the posterior distributions of neuronal model parameters—balances model complexity and accuracy in generative (i.e., forward) models of neuroimaging data. The open source platform of DCM allows researchers from diverse disciplines to formulate hypotheses and test them within a fairly standard framework. However, innovations are still required to improve group studies, hierarchical modelling of individual differences, and clinical applications.

We use dynamic causal modelling to invert a canonical microcircuit model of resting state MEG data in healthy adults (Friston et al., 2012, Friston et al., 2003). This entails the variational Bayesian inversion of biologically-informed forward models of neurophysiological observations, under the Laplace assumption; i.e., assuming Gaussian posterior probabilities over unknown parameters (Zeidman et al., 2022, Friston et al., 2007). In DCM, gradient optimisation of variational free energy is used for approximating the posterior probability density over unknown model parameters, and the model evidence (i.e. marginal likelihood) (Friston et al., 2007, Friston et al., 2008). The free energy provides a lower bound on log-model evidence, which represents the model accuracy adjusted for complexity. Model evidence associated with alternate hypotheses—about the underlying generators of data— are compared using Bayesian model reduction (BMR). This identifies the most likely explanation for the empirical data (Kass and Raftery, 1995, Friston and Penny, 2011). Crucially, BMR enables the computationally efficient evaluation of posteriors and model evidence under a reduced prior; i.e., a model specified in terms of new prior constraints (Friston et al., 2018, Friston and Penny, 2011). We estimate the parameters and evidence for a full model of a given dataset and then use BMR to evaluate posteriors and model evidences under alternative priors (Friston et al., 2018, Friston and Penny, 2011, Friston et al., 2019). At the second (e.g., group) level, hierarchical or parametric empirical Bayes (PEB) allows one to include empirical priors of interest (Friston et al., 2015, Friston et al., 2016). Usually, second level models apply priors that are conserved over multiple participants. This means the first level corresponds to a within-subject analysis, while the second level is a between-subject or group analysis (using subject specific posterior estimates as inferred by the first level). The combined hierarchical model can then be assessed in terms of its evidence, and subjected to BMR to test different hypothesis at the within or between-subject level.

In DCM, the prior mean and covariance of unknown parameters are specified to accommodate physiological interpretability and model stability, respectively. Effectively, (informative) priors provide constraints that enable the inversion of otherwise over-parameterised models (Friston et al., 2003). Using DCM, differences among individuals (or groups) can be characterised in terms of post hoc associations between connectivity parameters and clinical, pathological or cognitive measure of interest, or by differential model evidence, or conditional densities over models’ probabilities (Adams et al., 2021, Stephan et al., 2009, Rae et al., 2016, Passamonti et al., 2012). However, a more principled Bayesian methodology is to incorporate between-subject variables as priors on the generative model of brain physiology. Bayesian model reduction and group-DCM can then be used to test whether such (empirical) priors increase model evidence compared to conventional (or ‘flat’) priors. The source of empirical priors could be demographics (e.g. age), the burden of neuropathology (e.g. from PET scanning), or multi-modal measure (e.g. MRS). In this study, the DCM (at the first and group levels) leverages the spectroscopy information to impose constraints on synaptic physiology and assess their ‘goodness’ in terms of model evidence. If the embedding of empirical priors into a DCM leads to higher model evidence, one can infer the importance of the measured process in the generation of observed neurophysiology. We use MRS data, noting that GABA and glutamate changes are associated with many neurological conditions.

In this paper, we compare the evidence for different hypotheses about the association between MRS estimates of neurotransmitter concentration and synaptic function. See Figure 1 for an overview of the method. For a given set of connections, we use a hierarchical optimisation (a greedy search followed by a constrained optimisation) to identify the (linear or nonlinear) function of the MRS estimates—imposing prior constraints—that maximises model evidence at the group level. We then use model reduction to determine which subsets of synaptic parameters are sensitive to individual differences in neurotransmitter concentration. This approach complements previous work by (Stephan et al., 2009, Sokolov et al., 2019) who used diffusion-weighted imaging tractography as empirical priors on connectivity parameters that are inferred from functional magnetic resonance imaging time series. However, unlike previous work, we do not assume a one-to-one relation between MRS measures and connections in the generative model. Rather, we seek evidence to find which synaptic connections are informed by the empirical measures.

**Figure 1.**
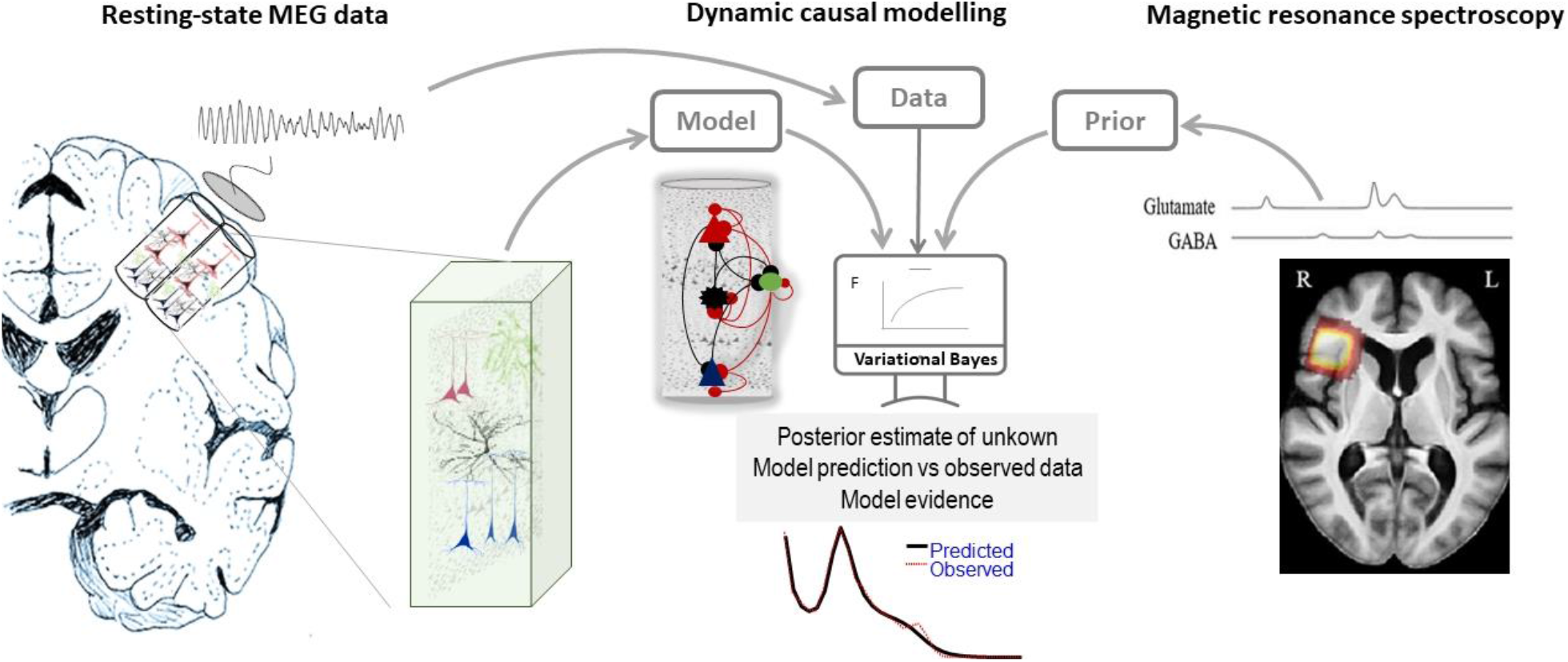
Dynamic causal modelling (DCM) can be informed by physiological or pathophysiological measures. DCM is the variational Bayesian inversion of neuronal models given neuroimaging Data: e.g., MEG. Given some data, DCM infers unknown synaptic parameters under a given (physically plausible) model and estimates the evidence for that model. The DCM parameter estimates can be informed by empirical priors; here, based on neurotransmitter concentration, as measured by magnetic resonance spectroscopy. Neurotransmitter concentration is proposed to modulate the synaptic efficacy of neuronal microcircuits that underlie neuronal dynamics that, in turn, generate electromagnetic signals. The comparison of such empirical priors — in terms of model evidence — extends Bayesian inferences about aspects of brain physiology which cannot be otherwise captured by non-invasive functional neuroimaging. The effect of the neurotransmitter concentration might manifest on one or more synaptic connections in the cortical microcircuit. By specifying which parameters are subject to empirical priors, one can then compare ensuing models to identify which kinds of synaptic connections are modulated by neurotransmitter concentrations. The MRS traces and heatmap are from (Murley et al., 2020) with permission.

In the following sections, we describe the multi-modal dataset (MEG and 7T-MRS) and the first-level DCM used to fit cross-spectral density data features. We then describe the second (group) level PEB model of the mapping between neurotransmitter concentration and synaptic physiology. We present the results of first-level and group-level DCMs, using MRS data as empirical priors. Finally, we discuss the potential applications and limitations of the method. A glossary and definitions of acronyms and variables used in this paper are provided in Tables 1 to 4.

**Table 1:** Acronyms

| Acronyms | Description |
| --- | --- |
| AMPA | α-amino-3-hydroxy-5-methyl-4-isoxazolepropionic acid |
| BMR | Bayesian model reduction |
| CMM | Conductance mesoscopic model |
| DCM | Dynamic causal modelling |
| FT | Fourier transform |
| fMRI | Functional magnetic resonance imaging |
| GABA | Gamma-aminobutyric acid |
| GLN | Glutamine |
| GLU | Glutamate |
| MEG | Magnetoencephalography |
| MRS | Magnetic resonance spectroscopy |
| NMDA | N-methyl-D-aspartate receptors |
| PEB | Parametric empirical Bayes |
| Odd/Even PSD | Power spectral density that was derived from odd or even trials in MEG data |
| RIFG | Right inferior frontal gyrus |
| sp, ss, in, dp | Superficial pyramidal cells, spiny stellate excitatory neurons, interneurons, deep pyramidal cells |
| $X$ | Design matrix of second level model (equation 6) with dimensions $n \times r$ ($n$ subjects and $r$ covariates) |
| $\theta^{(1)}$ | Vector of inferred parameters at first level DCM over all subjects with dimensions $np \times 1$ ($n$ number of subjects and $p$ number of inferred parameters in each DCM) |
| $\theta^{(2)}$ | Second level parameters with dimensions $r \times 1$ ($r$ number of covariates) |
| $\Gamma_{\varphi}$ | Scalar function of MRS data which is parametrised with a vector of parameters $\varphi$ |
| $\phi^1$ | Vector of inferred parameters at first level DCM from all subjects that are influenced by MRS data. |
| $\Pi^{(2)}$ | Precision matrix at the second level with dimensions $p \times p$ ($p$ number of parameters at the first level) |

## 2 Material and Methods

### 2.1 Participants

Eleven healthy adults (age 63-77 years, five women) participated after providing written informed consent. The study had ethical approval from the Cambridge 2 Research Ethics Committee. They were free from neurological or major psychiatric disorders and took no regular medication. Each participant underwent 7T MRI and MRS (Murley et al., 2020) and task free resting-state MEG data, on separate days.

#### 2.1.1 7T MRS data

Ultrahigh field magnetic resonance data were acquired using a Siemens Terra scanner at the Wolfson Brain Imaging Centre using short-echo semi-LASER sequence (repetition time/echo time = 5000/26 ms, 64 repetitions (Deelchand et al., 2015)), and VAPOR water suppression calibration (Gruetter and Tkáč, 2000). A 2cm isotropic cubic voxel was located over the right inferior frontal gyrus (RIFG). The MRS voxel was placed using conventional anatomical landmarks by the same operator (AGM) prior to each scan. Anatomical variants (such as absent diagonal branches of the Sylvian fissure) could in principle affect voxel placement accuracy. However, there was a very close overlap of the prefrontal MRS voxels across individuals (Figure 1 of (Murley et al., 2020)). Details of the voxel placement and MRS quality assurance metrics are provided with open access in Murley et al. (2020).

The details of the MRS processing have been reported previously in (Murley et al., 2020, Murley et al., 2022). In brief, the spectra were pre-processed to remove effects of eddy currents and frequency/phase shifts using MRspa (Dinesh Deelchand, University of Minnesota, www.cmrr.umn.edu/downloads/mrspa). The LCModel method (Version 6.2-3) was used to quantify glutamate and GABA between 0.5 and 4.2 ppm (Provencher, 1993). A MP2RAGE sequence (repetition time =4300 ms, echo time = 1.99 ms, resolution = 99 ms, bandwidth= 250 Hz/px, voxel size = 0.75 mm3, field of view =240240157 mm, acceleration factor (A>>P) = 3, flip-angle = 5/6 and inversion times = 840/2370 ms) was acquired for co-registration and segmentation, using SPM12 for partial volume correction, from fractions of grey matter, white matter and CSF.

The MRS voxel size was constant across participants (2cm isotropic). This voxel will contain varying partial tissue volumes, and derive from marginally different brain regions among individuals. However, we used a MRS sequence with outer voxel suppression pulses to optimise localisation and used regression analysis to correct for variation in age, sex and partial volume before modelling (see (Murley et al., 2020) for details). We used rigorous localisation standards, together with a generalized linear model to remove the effect of age, sex and correct for partial volume effects. Residual glutamate and GABA values were used for subsequent analysis (Murley et al., 2020). Note that ultrahigh field MRS (7T-MRS) can better distinguish between glutamate from within the Glx “peak”, and GABA, compared to high field MRS (3T-MRS).

#### 2.1.2 MEG data

Resting state MEG data were collected during two recordings, each of five minute duration, on a different day to the MRS, with eyes closed using an Elekta Vector View system with 204 planar gradiometers and 102 magnetometers. MEG data were recorded continuously with 1000 Hz sampling rate. Participants’ horizontal and vertical eye movements were recorded using bipolar electro-oculogram and electro-cardiogram electrodes. Five head position indicator coils were placed on an electroencephalography cap to track the head position. Three fiducial points (nasion, left and right pre-auricular) and a minimum of 100 head shape points were digitised using Polhemus digitization.

The data were pre-processed using the Elekta Neuromag toolbox (Elekta Oy), with MaxFilter v2.2.12 for detection and interpolation of bad sensors, and signal space separation to remove external noise from the data and correct for head movement correction. The data were then high-pass filtered at 1 Hz, stop-band filtered around [22 to 24] Hz and [47 to 51] Hz and divided into epochs of one second duration. We used the Field trip Toolbox (Oostenveld et al., 2011) for detection and removal of eye movement artefacts and discontinuities.

We applied empirical Bayesian inversion in SPM12 for source inversion and extraction of the right IFG source time series for subsequent analyses (Litvak et al., 2011). We concatenated the data of the two recordings and then divided the source data into two sets, one comprising the odd numbered epochs (‘odd data’) and one with the even numbered epochs (‘even data’) for each participant to assess the reliability of the results (Litvak et al., 2015). We separately averaged power spectral responses of the odd and even datasets. The two power spectral densities (referred to as Odd PSDs and Even PSDs) are used as the data features in the DCM of cross-spectral density.

### 2.2 First level analysis using dynamic causal modelling of resting states MEG data

To infer the neurophysiological parameters generating observed resting state MEG data, we used “DCM for cross spectral density” in SPM12 (Friston et al., 2012, Moran et al., 2007, Moran et al., 2011). Spectral features in the resting state MEG were used to infer synpatic parameters of a biophysically-informed neuronal mass model, together with model evidence. DCM for cross spectral density assumes that recorded electrophysiological oscillations are due to finite responses of neuronal dynamics, under endogenous random fluctuations (Basar et al., 2012, Haken, 1977).

We used a conductance based neuronal mass model as shown in Figure 2 (known as “CMM_NMDA” model in SPM12) (Moran, 2015, Moran et al., 2013, Shaw et al., 2017). The conductance based model represents the actvity of cortical columns based on the interactions of four neuronal populations: excitatory spiny stellate cells, superfical pyramidal cells, inhibitory interneurons, and deep pryramidal cells as shown in Figure 2.

**Figure 2.**
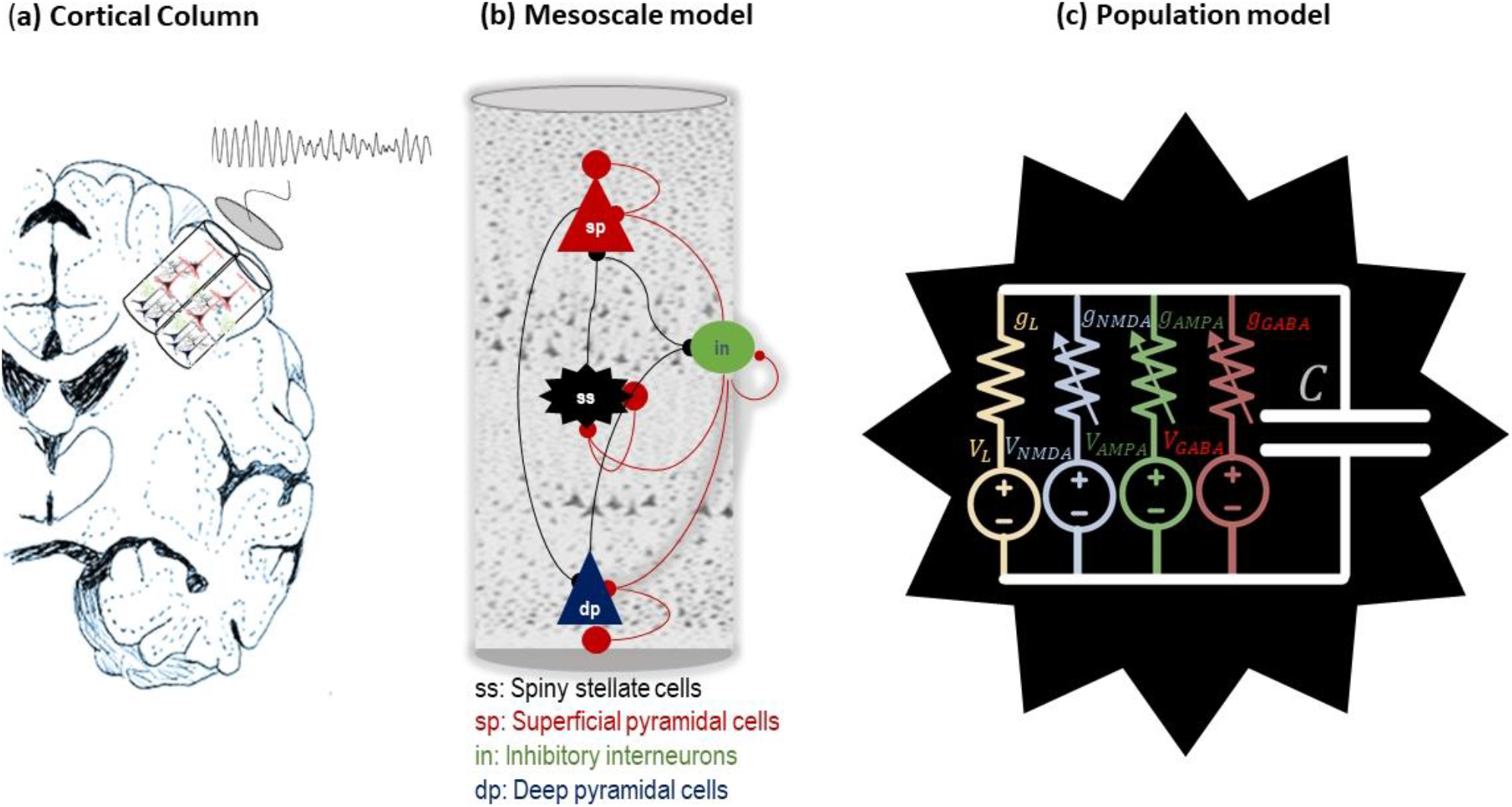
The conductance based neural model. Panel (a) shows a cartoon of a cortical column which generates electrical brain activity that can be captured by neuroimaging modalities such as MEG. Panel (b) illustrates the neuronal model which is divided into three layers where superficial (sp) and deep pyramidal (dp) cells are in the top and bottom layers, respectively, excitatory interneurons (spiny stellate cells, ss) are located in layer four, and inhibitory interneurons are distributed across all layers and are modelled using one population that interacts with all other populations. In addition, each population is equipped with a self-inhibition which assures dynamical stability around a stable fixed point. Panel (c) illustrates the population model, each neuronal population is governed by the Morris-Lecar model. This model explains the dynamics of different ion currents: NMDA, AMAP and GABA and passive ion current and membrane capacitance as explained in equation 1. Brain, Resistor, capacitor and voltage icons by Michael Senkow from thenounproject.com, CC BY 3.

Each population is represented by a Morris–Lecar model (Moran et al., 2013). The dynamic of each population is govened by stochastic differential equations that emulate the dynamics of pre/post-synaptic potentials, firing rates and membrane conductances. In a typical neruonal population, the dynamics of membrain potentials, *V*, and conductances of an ion channel, *g*_*_, are govered by the following equations:

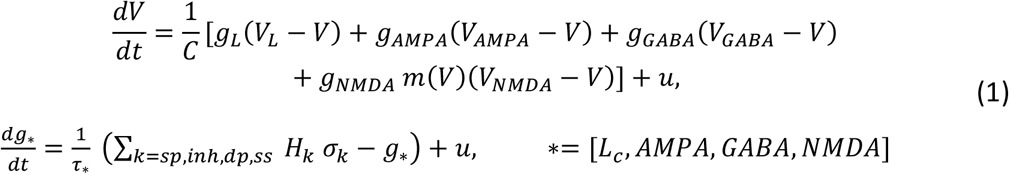

In equation 1, *C* is the membrane capacitance, *u* is random endogenous fluctuation, *L_c_* is passive leak current where its conductance is constant, *g*_*_ is conductance associated with ion channels/receptors with time constant *τ*_*_, *V_AMPA,GABA,L,NMAD_* are the reversal equilibrium potentials of the ion channels. The *m*(*V*) in equation 1 is the actvity-dependent magnesimum channels which is given by 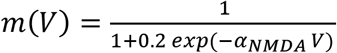. The term *σ*_*k*_ in the equation 1 is the non-negative afferent presynaptic firing from population *k*, which is scaled by afferent intrinsic connectvity *H*_*k*_.

The scaled presynaptic firing rates are a proxy for neurotransmitter levels measured by MRS. Therefore, we hypothesise a relation between the MRS glutamate and GABA measures and excitatory and inhibitory *H*_*k*_ connections, which scale presynaptic actvity. In other words, the MRS glutamate and GABA can be used as empirical priors on excitatory/inhibitory connections in the generative model.

Mathematically the temporal dynamics of the conductance based model can be written in the canonical form of a dynamical system as follows (Friston et al., 2012):

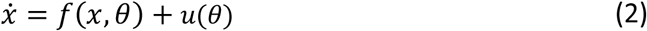

Where *θ* denotes a vector of all unknown parameters (i.e., *H*_*k*_,*τ*_*k*_), *x* is the state of neuronal populations in the model, *f*(*x, θ*) is a function that is the concatenated version of the right hand sides of equation 1—over all populations—and *u* represent endogenous fluctuations, conventionally modelled by (structured) pink noise. The noise in equation 2 has a cross-spectrum *g*_*u*_(*ω, θ*) = *FT*(*E*[*u*(*t*), *u*(*t* − *τ*)]): Please see Table 2.

**Table 2:** Glossary of variables and expressions in conductance-based neuronal model.

| Variable | Description |
| --- | --- |
| $u$ | Exogenous input (scalar) to membrane equation and conductance equation within each population. |
| $V$ | Mean depolarisation of a neuronal population. It is a scalar variable per each population. |
| $\sigma(v)$ | The neuronal firing rate (scalar function) – a sigmoid squashing function of depolarisation |
| $L$ | Lead field vector mapping from (neuronal) states to measured (electrophysiological) responses |
| $g_x(\omega), g_o(\omega), g_y(\omega)$ | Spectral density of (neuronal) state fluctuations, observation noise and measurement, respectively. These are vector valued functions. |
| $\nabla_x f$ | System Jacobian or derivative of system flow with respect to (neuronal) states (matrix valued function). |
| $k(t) = FT[K(\omega)]$ | First order kernel mapping from inputs to responses; c.f., an impulse response function of time. This is the Fourier transform of the transfer function. |
| $K(\omega) = FT[k(t)]$ | Transfer function of frequency, modulating the power of endogenous neuronal fluctuations to produce a (cross spectral density) response. This is the Fourier transform of the kernel. |

One can approximate the dynamics of equation (2) with the (first order) linearised model 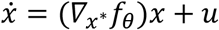 denotes the Jacobian at *x*^*^) with the spectral response:

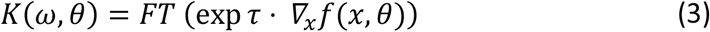

Given the spectral response of the system, the generative model of neuronal activity, *g*_*x*_(*ω*), can be calculated as follows:

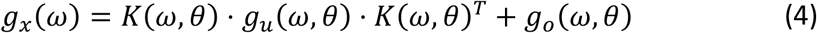

In equation 4, *g*_*o*_(*ω, θ*) represents the spectrum of the observation noise, which is a sum of common and source specific noise. The spectral response in sensor space can also be generated by inclusion of a forward electromagnetic model into equation 4, which is denoted by *L. M*(*ω*) (*L* is the gain and *M* is the head model), as follows:

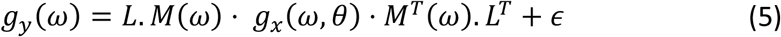

In equation 5, *g*_*y*_(*ω*) is the cross spectra of the MEG data and *ϵ*∼*N*(0, *σ*) is a random effect (with unkown covariance). Because we perform the DCM analysis in source space, the forward electromagnetic model in this case just applies a scaling parameter. In the DCM, we do not pre-define which neuronal population(s) contribute to MEG data. Instead, we estimated the degree to which each population (e.g. superficial, deep pyramidal or inhibitory population) contributed to the generation of the MEG data (see Pereira et al 2021 for details specific to conductance based models). Recently clinical and interventional applications support this approach (Adams et al., 2021, Shaw et al., 2021, Gilbert et al., 2016, Symmonds et al., 2018).

The unkown parameters in the DCM are specified as log-scale values. This means that the parameter vector in DCM is a random variable, which is given by 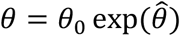. Here, *θ*_0_ is a biologically informed scaling for the parameter and 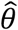is a random variable, with prior normal density 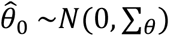 of zero mean and covariance ∑_*θ*_. This expresses the belief about the range over which parameters can vary. Note that scale parameters of this sort cannot be less than zero; for example, distances and lengths or rate and time constants or precisions and variances.

### 2.3 Second level analysis: group DCMs informed by empirical priors

In many translational neuroscience studies, the goal is to test hypotheses at the group level, where subject-specific information is given by summary statistics. In PEB, this subject-specific information is the posterior over key model parameters at the first level (Zeidman et al., 2019a, Zeidman et al., 2019b, Friston et al., 2015, Friston et al., 2016). At the second level the objective is to test whether synaptic connections depend on between subject variables, such as age, disease-severity, genetics, diffusion tensor imaging and MRS (Friston et al., 2016). In other words, PEB seeks to explain intersubject variability on one or more first level model parameters.

Mathematically, we denote a vector of model parameters at the first level DCM, over all participants, by a column vector *θ*^(1)^ (superscript ‘1’ denotes the first level analysis) with dimension *np* × 1 (*n* number of participants and *p* is the number of parameters for each participant). Then the generative model at the second level (i.e., random effects on the parameters) is given by (Friston et al., 2016):

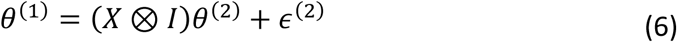

Where *X* ∈ *R*^*n*×*r*^ is the design matrix with *r* ≥ 1 covariant (the first regressor of X is always equal to one and reflects the group mean) and a column vector *θ*^(2)^ (superscript ‘2’ denotes the second level) contains the second level parameters. The symbol ⊗ is the Kronecker product and *I* is the *p* × *p* identity matrix. The random effects have a Gaussian distribution *ϵ*^(2)^∼*N*(0, *Π*^(2)^) (where *Π*^(2)^ is precision matrix or inverse of the covariance). The precision matrix is parameterised with a single (hyper-precision) parameter, γ, as follows (Friston et al., 2016):

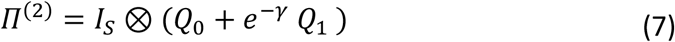

In equation 7, *Q*_0_ ∈ *R*^*p*×*p*^ is the lower bound on the precision, defined with a small positive value). The (hyper)parameter, *γ*, scales a precision matrix *Q*_1_ ∈ *R*^*p*×*p*^, which is (by default) 16 times the prior precision of the group mean (Zeidman et al., 2019b): this hyperprior ensures that random effects are small in relation to prior uncertainty about the parameter in question.

The critical question is whether an alternative prior increases or decreases model evidence (Friston et al., 2016). This is an optimisation problem, where the objective function is the free energy of the PEB model. Inversion of the hierarchical model is computationally expensive if all parameters of the first level need to be inferred every time the second level parameters change (Raman et al., 2016). However, BMR can be used to re-evaluate first level posteriors under updated second level parameters (Friston et al., 2016, Litvak et al., 2015). This significantly improves the efficiency of the inversion for PEB models.

### 2.4 Investigating relation between MRS and synaptic connections

#### 2.4.1 Problem setting

We test whether excitatory and inhibitory synaptic connections depend on (i.e., are functions of) glutamate and GABA measures, respectively. We used PEB to specify and compare the evidence for different functions of MRS measures, as illustrated in Figure 3.

**Figure 3.**
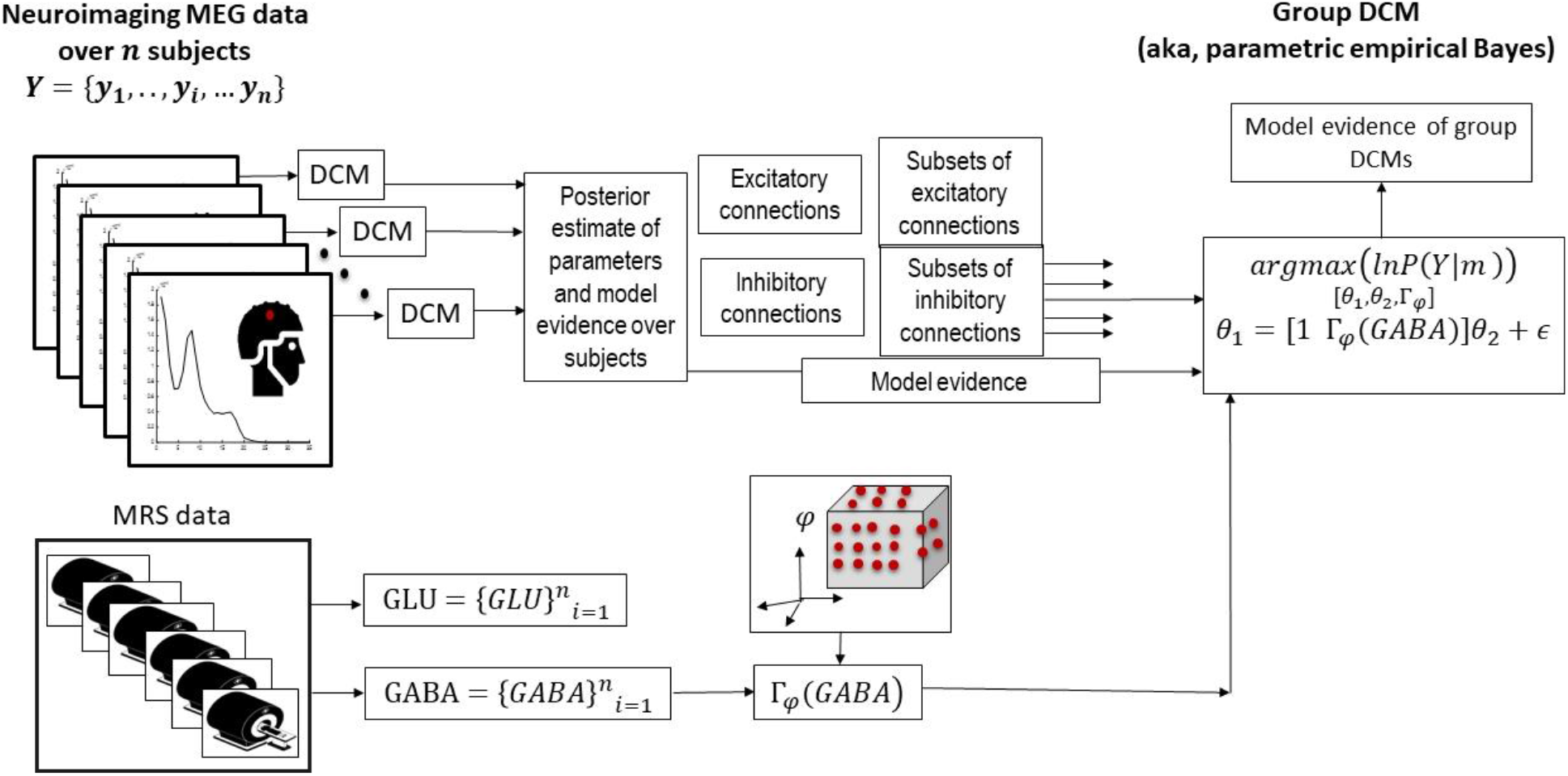
MRS informed group DCM for MEG. This graphic illustrates a group DCM (aka PEB model) inversion using MRS data as empirical priors. First level DCM (left side of panel) infers synaptic physiology from each participants’ data. A function of MRS glutamate or GABA measures (bottom panel) are considered as empirical priors for certain parameters in the group DCM (right panel). Bayesian model reduction evaluates the effect of empirical priors and evaluates the model evidence for the group DCM. The objective is to find the optimal function of MRS measures—that inform synaptic parameters in the CMM-NMDA model—by maximising PEB model evidence. Image credit for the MRI scanner icon by Grant Fisher, TN from thenounproject.com, CC BY 3.

To identify which synaptic parameters were sensitive to neurotransmitter levels, we grouped the inferred synaptic connections into excitatory and inhibitory subsets. In detail, for the *l*_*ex*_ (*l*_*Inh*_) excitatory (inhibitory) connections in the neuronal model, we define 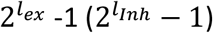 different possible combinations. The ensuing subsets of the connections (over all participants) are considered as the dependent variable of the PEB model (i.e., the right-hand side of equation 1). This model encodes a hypothesis about the relationship between (each subset of) connections, *ϕ*^(1)^ = {*ϕi*}^*k*^_*i*=1_ and a particular function of MRS data: Γ_*φ*_(*MRS*) (where Γ is a smooth and monotonic function with unknown parameters, *φ*), which can be expressed as follows:

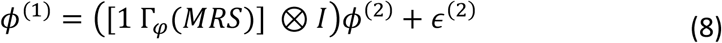

In equation 8, there are two sets of unknowns that need to be inferred; namely, *(i)* the subset of connections *ϕ*^(1,2)^ (superscript 1 and 2 denotes to first and second level parameters, respectively) and *(ii*) the function or map Γ_*φ*_(*MRS*). These unknowns are the hyper-parameters in equation 6. To find optimal hyperparameters, we recursively select subsets of synaptic connections and estimate the MRS function parameters, such that the model evidence in equation 6 is maximised. In other words, for any combination of synaptic parameters, we seek the MRS-informed PEB model with the highest evidence, using BMR. This allows us to identify the most likely solution (in terms of model evidence) from the model space tested (a set of potential monotonic relationships), to identify which set of synaptic parameters are informed by MRS data.

To cross-validate the results, we separately tested the relationship between inferred synaptic parameters and MRS by splitting the MEG data into odd and even numbered epochs (odd-Data and even-Data). The ensuing procedure is shown in Figure 4.

**Figure 4.**
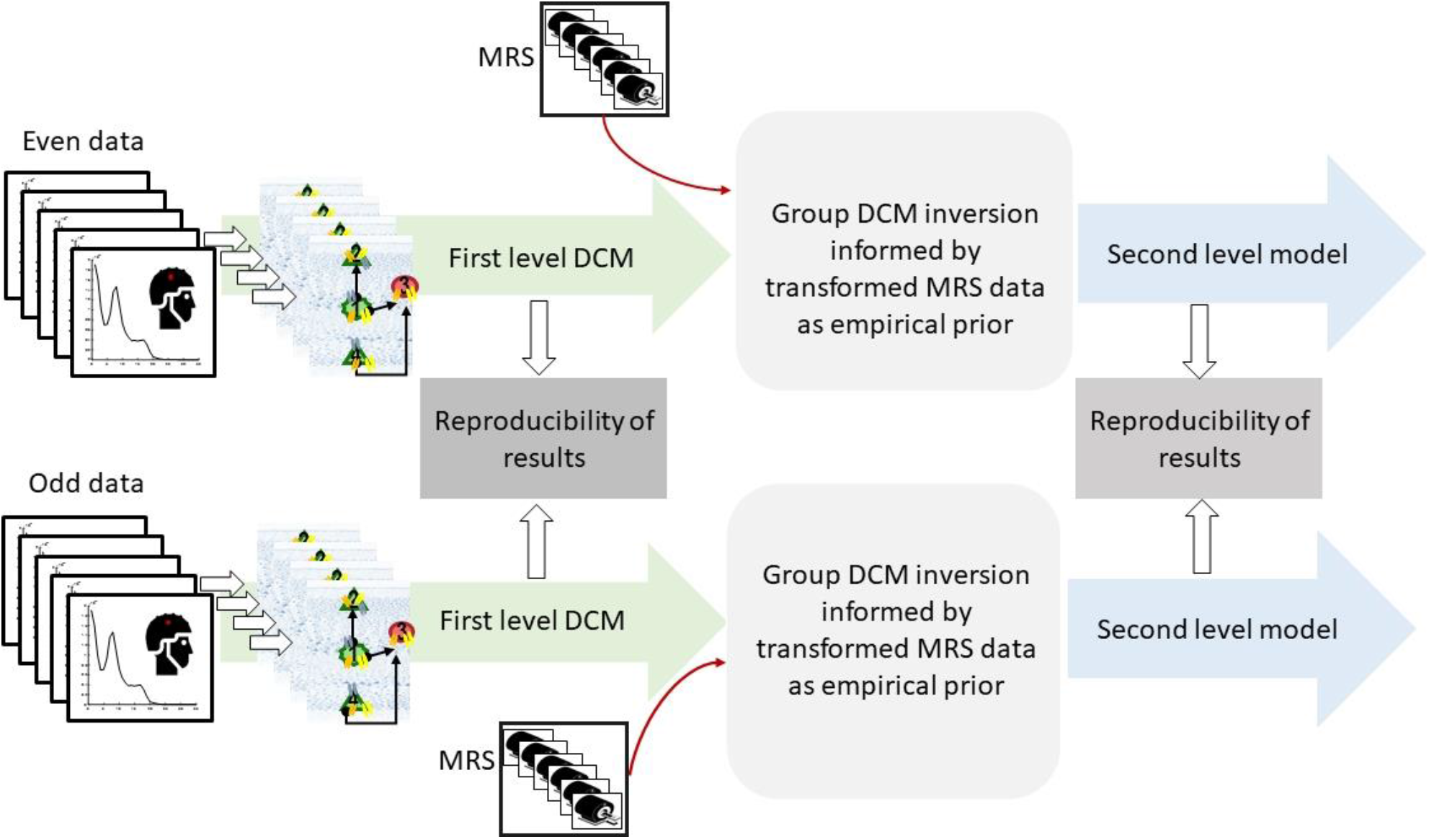
Reproducibility-check procedure. The MEG data for each participant is divided into two separate datasets, ‘odd data’ and ‘even data’. DCM for MEG is used to infer synaptic parameters for each. A post hoc correlation analysis is used to test agreement between the ensuing estimates. Parameter estimates that are consistent over the datasets are considered for further analysis. We perform the group-level inversion informed by MRS data for odd and even datasets on the selected parameters and explore which sets of connections are best informed by a function of MRS data. Finally, we compare the outcome of the PEB inversion and check the consistency of the results between the odd-data and even-data. MRI scanner icon by Grant Fisher, TN from thenounproject.com, CC BY 3.

#### 2.4.2 Functional form of the empirical MRS priors

The functional form of the MRS mapping is not known *a priori*. We therefore limit the search space over the MRS transformations to continuous and monotonic polynomial maps and sigmoid nonlinear functions, which cover a wide range of linear and nonlinear forms. The class of polynomials provides an approximation (Bishop, 2006) to any nonlinear monotonic form (Spivak, 2020):

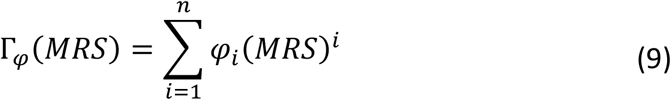

Here, *φ*_*i*_ are *n* unknown hyperparameters that need to be inferred by finding the values that maximise model evidence. This class of functions express different nonlinear relations including saturations (Stephan et al., 2009, Stefanovski et al., 2019), *n* = 1 implies a linear transformation of the MRS data.

Animal experiments suggest that changes in GABA and glutamate concentrations are related to neuronal responses via a sigmoid relationship (Benardete and Kriegstein, 2002, Dyke et al., 2017, Chebib et al., 2009). The general form of a sigmoid nonlinearity can be parameterised as follows:

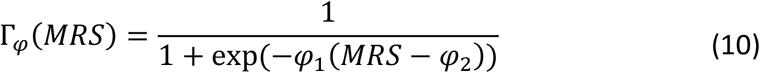

Here, the unknown hyperparameters *φ*_1_, *φ*_2_ are the slope and threshold parameters of the sigmoid transformation, respectively.

#### 2.4.3 Hyperparameter estimation

To estimate unknown hyperparameters, we first sorted the MRS measures from small to large values, which defined an interval of the real line (called the domain of the MRS data). We then examined the variation of the parameters under the MRS transformation. In the case of the polynomial form, we checked the monotonicity of the transformation. More formally, we denoted the space of monotonic functions by ℵ, and defined the function γ over the ordered domain set of the MRS data to identify the range for the variation of the parameters in the polynomial function. This function has the following properties:

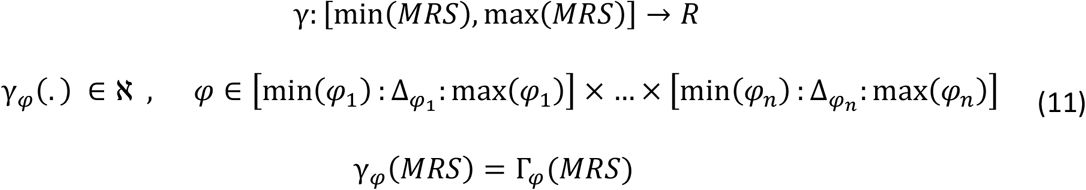

Each parameter in the *γ* map, *φ*_*i*_, varies with resolution (or step-size) 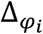. Note that *γ*_*φ*_ and Γ_*φ*_ have the same functional forms but with continuous and discretised domains. Having established the appropriate ranges (see Table 4), we used a hierarchical scheme to estimate the parameters of the map, Γ_*φ*_(*MRS*). For each set of synaptic connections, we calculated the model evidence—using BMR—for different values of the hyperparameters over the interval 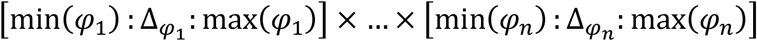. This constitutes an exhaustive grid search. This was followed by a constrained Newtonian search, within the neighbourhood of the selected grid point, to identify the precise value of the hyperparameters that maximised model evidence. We repeated this hyperparameter optimisation over all possible subsets of the synaptic connections and selected the combinations with the greatest model evidence.

**Table 3:** Parameters of the neuronal model (see also equation 1 and Figure 2)

|  | Description | Parameterisation | Prior |
| --- | --- | --- | --- |
| $\tau$ | Rate constant of ion channels | $\exp(\theta_\tau) \cdot \tau$<br>$\tau = [256, 128, 16, 32]$ | $p(\theta_\tau) = N(0, 1/16)$ |
| C | Membrane capacitance | $\exp(\theta_c) \cdot C$<br>$C = [128 \ 128 \ 256 \ 32]/1000$ | $p(\theta_c) = N(0, 1/16)$ |
| H | Intrinsic connections | $\exp(\theta_H) \cdot H$<br>$H = \begin{bmatrix} 8 & 0 & 2 & 0 \\ 4 & 8 & 8 & 0 \\ 0 & 0 & 32 & 128 \end{bmatrix}$ | $p(\theta_{Ha}) = N(0, 1/32)$ |

**Table 4:**
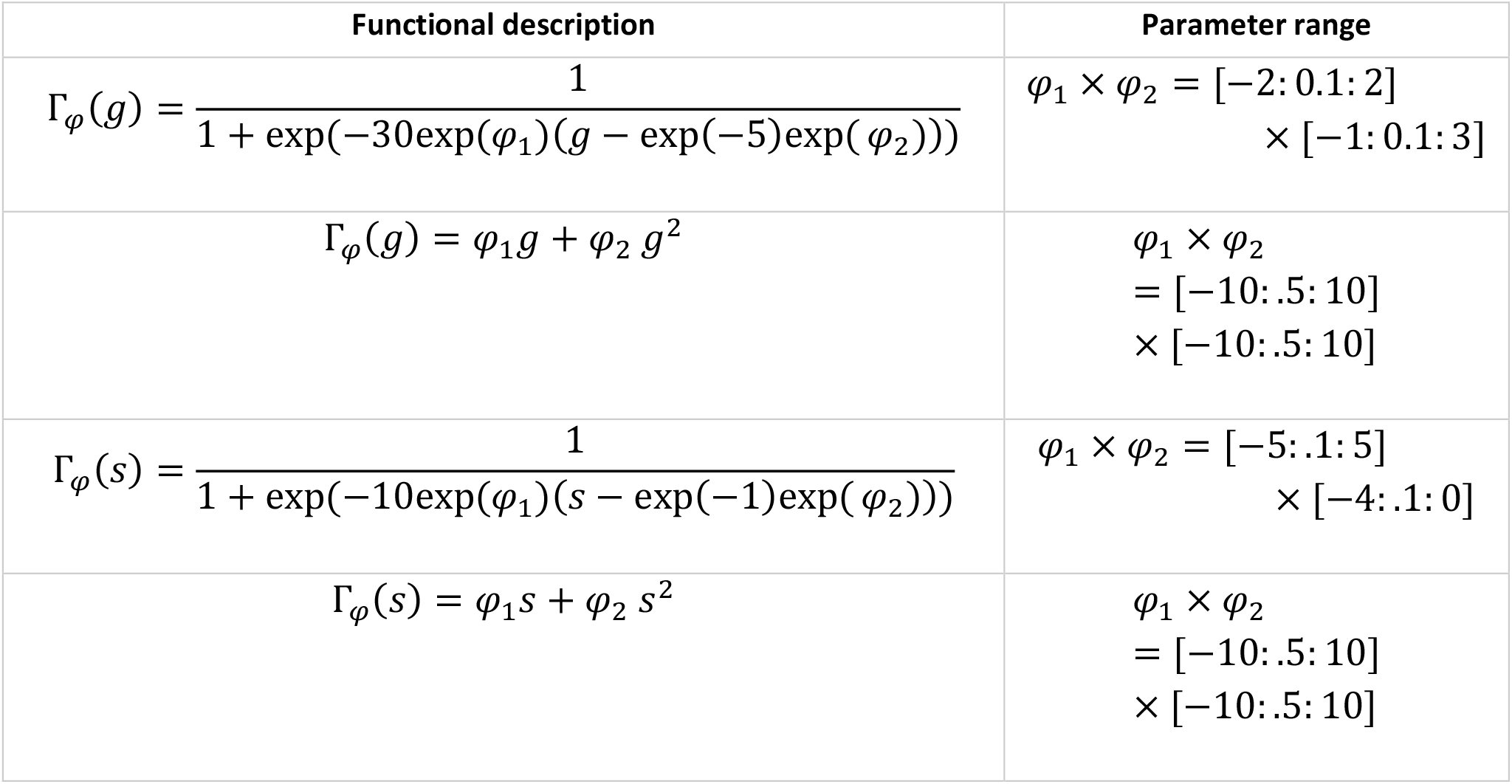
Parameterisation of MRS GABA (variable g) and glutamate (variable s) transforms.

| Functional description | Parameter range |
| --- | --- |
| $\Gamma_{\varphi}(g) = \frac{1}{1 + \exp(-30\exp(\varphi_1)(g - \exp(-5)\exp(\varphi_2)))}$ | $\varphi_1 \times \varphi_2 = [-2: 0.1: 2]$<br>$\times [-1: 0.1: 3]$ |
| $\Gamma_{\varphi}(g) = \varphi_1 g + \varphi_2 g^2$ | $\varphi_1 \times \varphi_2$<br>$= [-10: .5: 10]$<br>$\times [-10: .5: 10]$ |
| $\Gamma_{\varphi}(s) = \frac{1}{1 + \exp(-10\exp(\varphi_1)(s - \exp(-1)\exp(\varphi_2)))}$ | $\varphi_1 \times \varphi_2 = [-5: .1: 5]$<br>$\times [-4: .1: 0]$ |
| $\Gamma_{\varphi}(s) = \varphi_1 s + \varphi_2 s^2$ | $\varphi_1 \times \varphi_2$<br>$= [-10: .5: 10]$<br>$\times [-10: .5: 10]$ |

## 3 Results

### 3.1 DCM for MEG

The spectral activity from both data sets per participant (even and odd data) are used as the data features for a conventional “DCM cross spectral density” to infer the parameters of the conductance based canonical microcircuit. The predicted and observed data are provided in the supplementary material Figure 13. Over all subjects, the mean variance of observed CSDs explained by the predicted CSD was 98%. The comparison of predicted and observed spectral data shows that the synaptic parameter estimates can replicate the spectral patterns of both even and odd data; with an example shown in Figure 5 (see the supplementary material for the remaining comparisons). The correlation between the free energies associated with DCM inversions of odd/even PSD data is shown in Figure 5.

**Figure 5.**
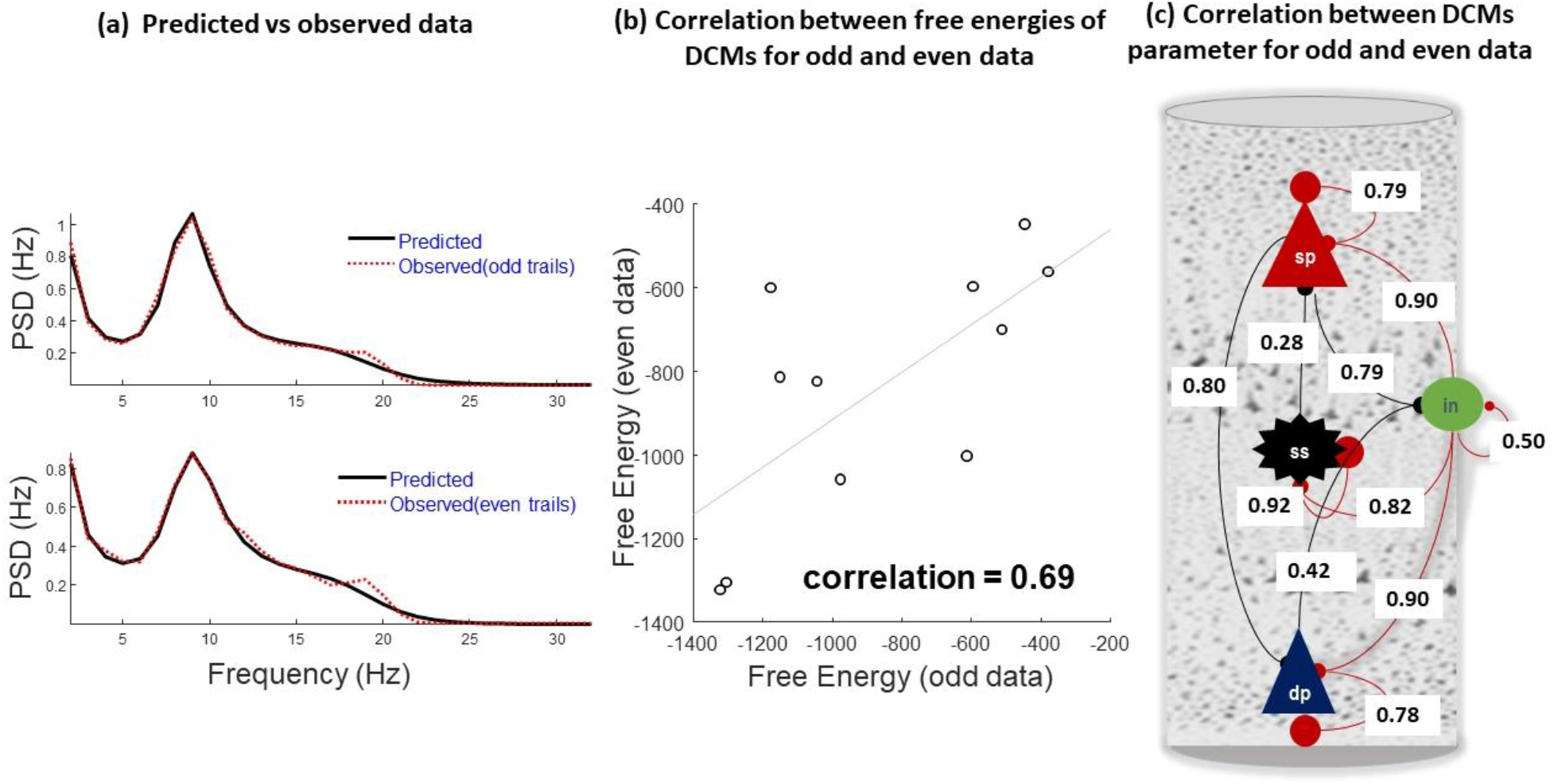
First level DCM results. Panel (a) shows predicted vs observed MEG data for odd and even data from a single participant. This panel shows that DCM can explain the spectral dynamics of each dataset well. Inversion results of the rest of the subjects are provided in the supplementary material, Figure 13. Panel (b) shows the correlation between free energies of odd and even datasets over participants (points in the plot). This correlation shows a moderately good level of agreement between the odd and even data across all participants. Panel (c) illustrates correlations between inferred synaptic connection between odd and even epochs. The inhibitory connections are shown in red and excitatory connections are shown in black.

To confirm the predictive validity of DCM, we calculated the correlation between the inferred synaptic parameters from odd and even PSDs data sets: see Figure 5. Most parameters were estimated reliably (figure 5). There were exceptions, such as the excitatory connections of the deep pyramidal and inhibitory connections and superficial pyramidal cells and inhibitory cells. The low correlation between estimates of some parameters is not surprising for complex nonlinear models with conditional dependencies among parameter estimates (Litvak et al., 2019). It implies that different combinations of synaptic parameters may generate similar physiological data. This means hypotheses may therefore be better tested in terms of model comparison, rather than focussing on maximum, *a posteriori* parameter estimates of single parameters (Rowe et al., 2010). In the following section, we use the reliable connections— with a correlation of greater than one half—to examine the effect of neurotransmitter concentrations as measured by MRS.

### 3.2 Associations between synaptic parameters and MRS measures

The most likely relationship between the synaptic parameters and neurotransmitter levels was identified through the hyperparameter optimisation.

#### 3.2.1 MRS-GABA data as an empirical prior on inhibitory synaptic parameters

MRS GABA could be evaluated as an empirical prior constraint on 127 combinations of inhibitory connections (2^7^-1, combinations of seven inhibitory connections in the conductance based canonical microcircuit model). As each combination might be constrained by empirical MRS GABA priors, Bayesian model reduction can be used to assess which of the 127 combinations and linear/nonlinear transformations of MRS GABA measures maximise model evidence. Alternatively, one can group certain synaptic parameters based on their common features (self-inhibition vs inter-regional connections or superficial vs deep connections), to identify the winning model ‘family’ for each subgroup of connections.

We grouped the inhibitory connections into four self-connections and three inter-lamina connections. By grouping the parameters in this way, we ask whether MRS GABA influences recurrent intra-laminar connectivity, or inhibition between layered populations. We assess the evidence for GABA effects, as mediated through second order polynomial and sigmoid transformations of the MRS measures. We systematically varied the hyperparameters of the sigmoid function for each subset of self-inhibitory synaptic connections and compared the resulting model evidence.

Figure 6 shows the consistency of the resulting free energy over odd and even epoch datasets. The correlation plot between the free energies of odd and even data models shows that the ranking of model evidence is consistent, where each ‘model’ is a hypothesis about how MRS data supplies prior constraints on synaptic parameters. Such agreement is an important validation of hypothesis testing, based on selecting models with the greatest evidence. Using PEB to make inferences at the between-subject level inherits this validity. The high reliability suggests that both group inversions converge to the same (global) minima. For each subset of self-inhibitory connections, the sigmoid MRS GABA hyperpriors maximised the model evidence. The winning models for each subset were then examined to determine which MRS mapping is most likely for each subset of self-inhibitory connections. As shown in Figure 7, priors using a sigmoid transformation of GABA concentration provided the most likely account of intersubject variation in synaptic connectivity; specifically, the recurrent connections or self-inhibition of superficial and deep pyramidal cells.

**Figure 6.**
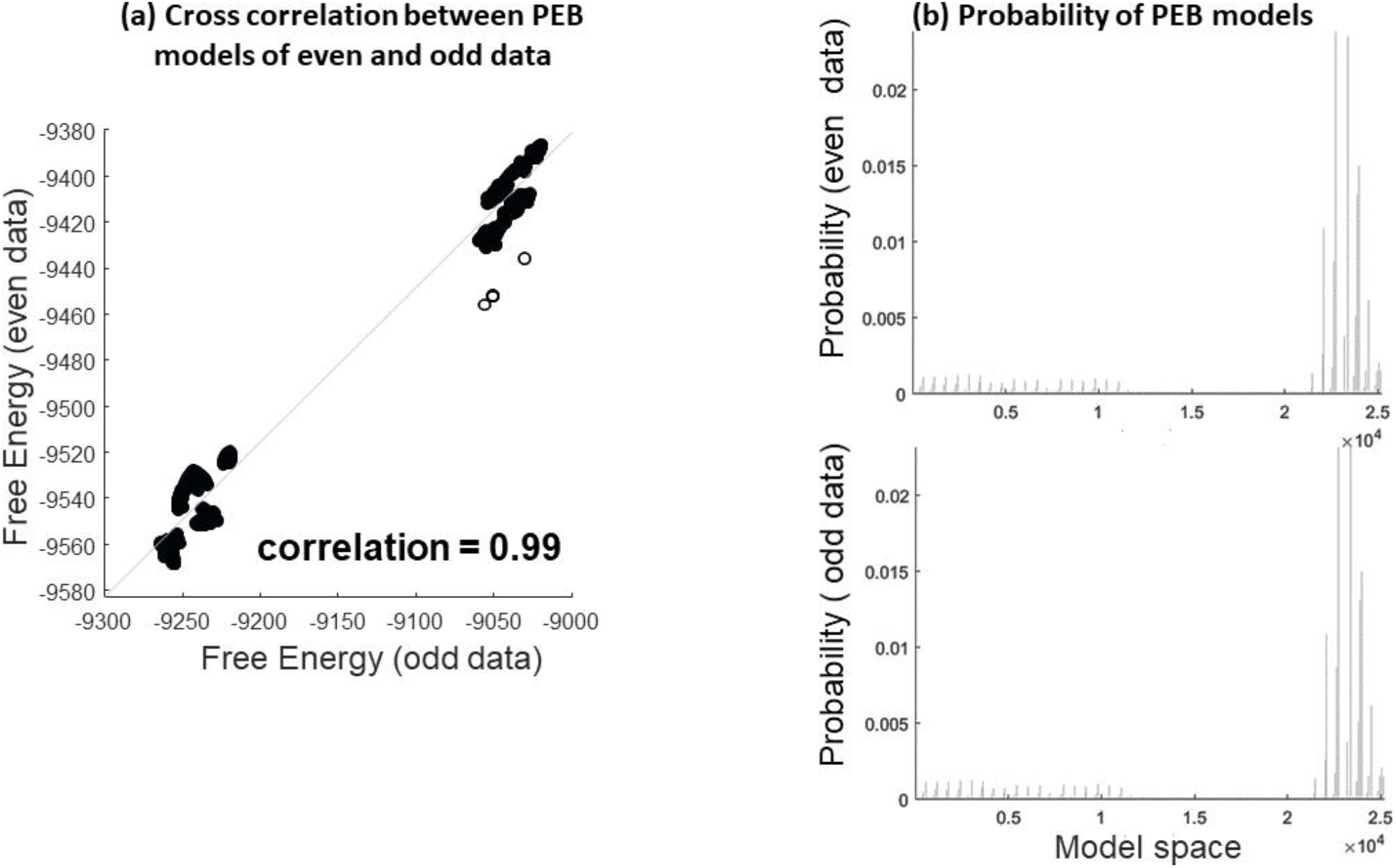
Correlation analysis, comparing the free energy of the PEB model over all subsets of self-inhibitory connections and all sigmoid transformations of GABA in odd and even data. This plot shows that the PEB approach consistently estimates the marginal likelihood of models in even and odd data. (b) The probability of different PEB models over model space. Both results show the same maximum. Due to having a large model space, the probability of models and the effect sizes (free energy) are small (this is known as model dilution).

**Figure 7.**
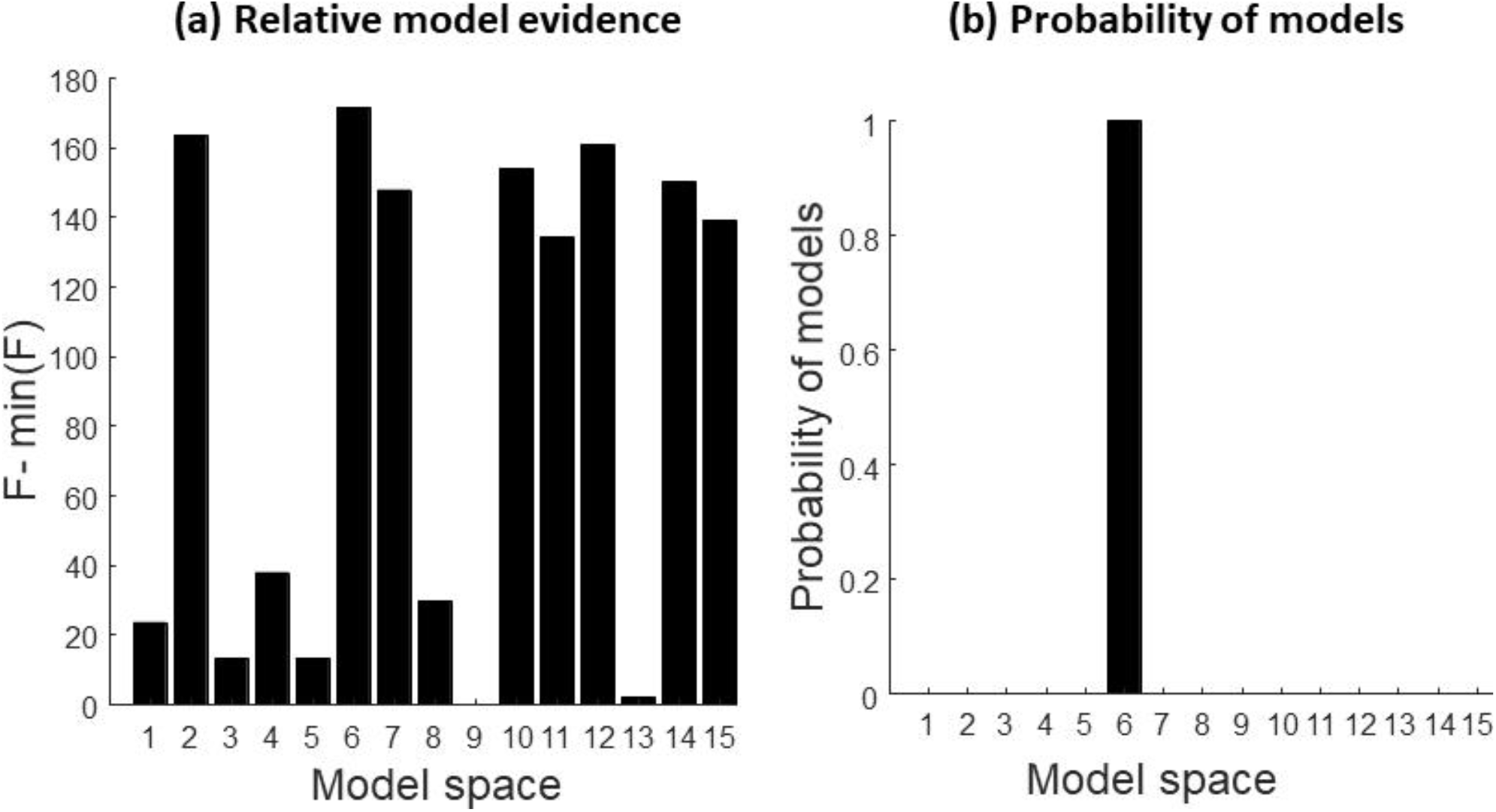
(a) The relative free energy of each subset of the self-inhibitory connections. (b) the probability of each model. The most likely “winning” model is 6 in which a prior of the sigmoid transformation of MRS GABA informs connections self-inhibition of superficial and deep pyramidal cells. Please see supplementary Figure 15 for definition of models in this graphic.

We repeated the same procedure with inter-laminar (i.e., intrinsic, between population) connections. The results are shown in Figures 8 and 9. MRS GABA provided an informative empirical prior for all inhibitory connections. The MRS transforms for odd and even datasets differ, but are similar in their functional form.

**Figure 8.**
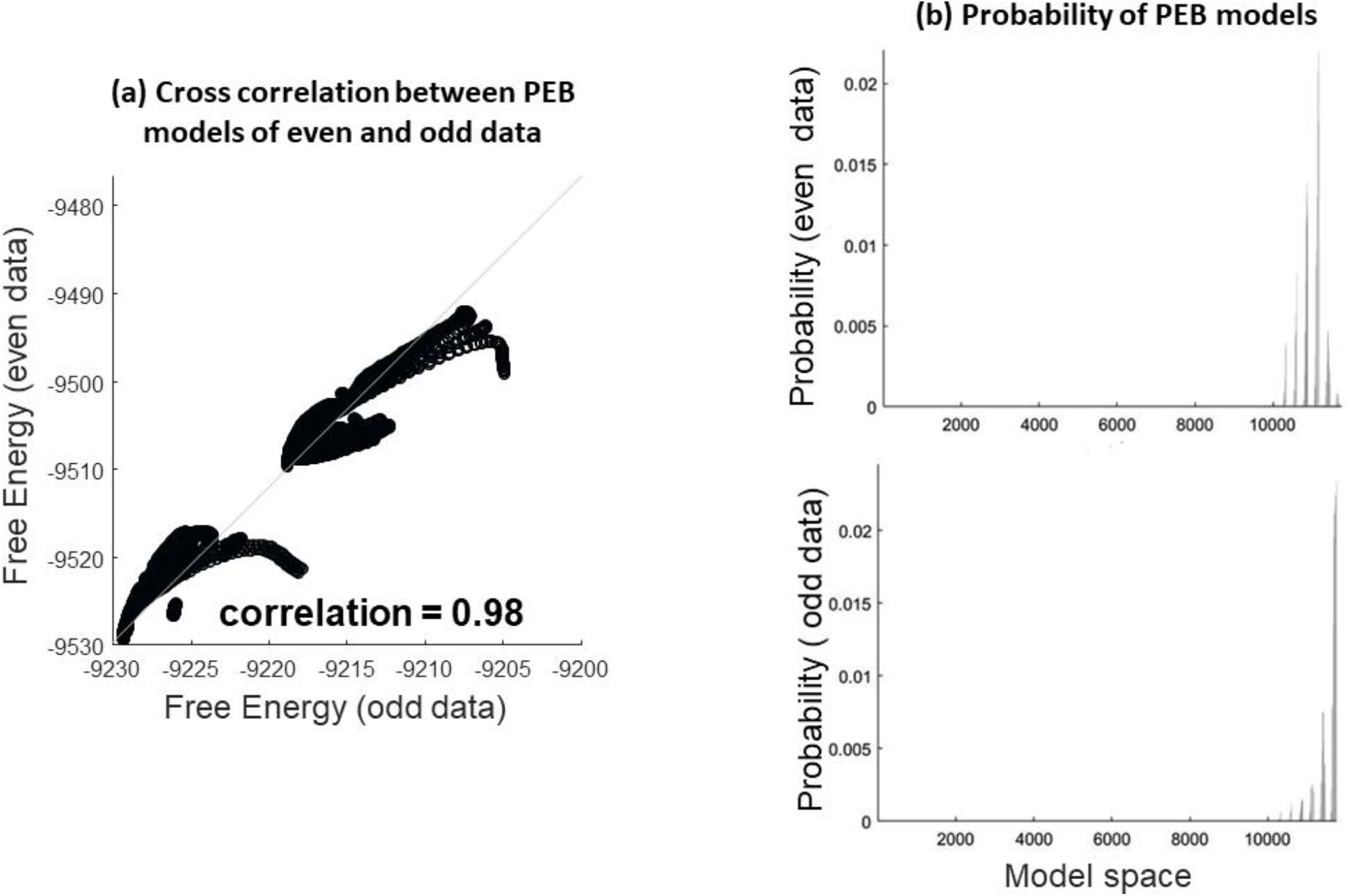
(a) This panel shows a correlation analysis between the free energy from the PEB model that encodes the relationship between all possible inter-regional inhibitory synaptic connections and all possible forms of sigmoid transformation of the MRS-GABA. These results suggests that the PEB approach reliably estimates model evidence in even and odd data. (b) The probability of different PEB models over model space. The maxima in the probability plots are not consistent due to the fact that there are two different MRS transformations with the greatest evidence for the odd and even data. The maximum probability in each plot is associated with the same combination of inhibitory connections.

**Figure 9.**
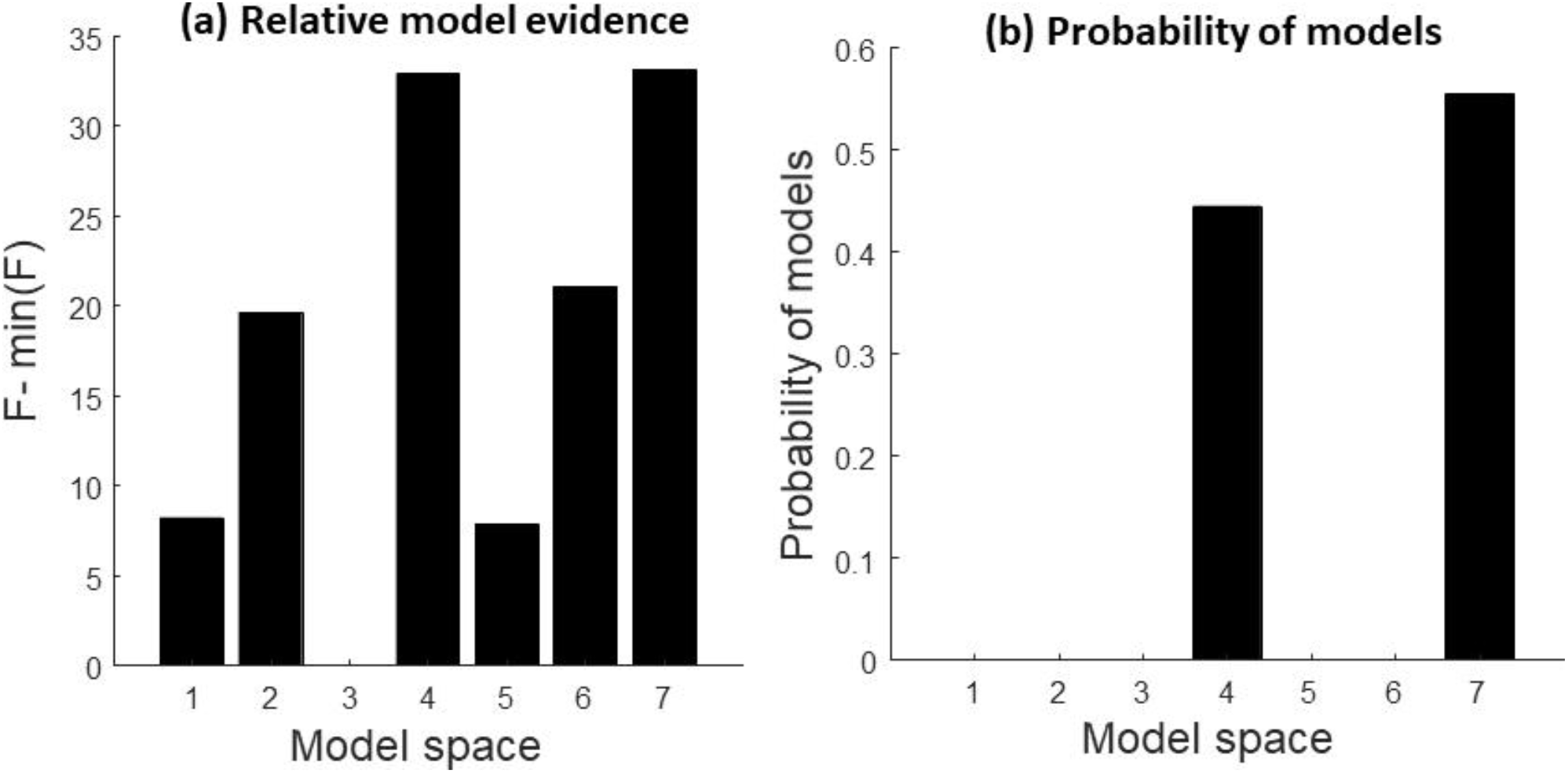
(a) The relative free energy for each subset of inter-laminar inhibitory connections for an optimal transformation is shown (b) The probability of each subset of synaptic connections is illustrated. The winning model identifies that a sigmoid transformation of MRS-GABA measures is likely to inform all inter-laminar inhibitory connections. Please see supplementary Figure 15 for definition of models in this graphic.

We repeated this procedure using a second order polynomial (the first order polynomial is contained within this model space) as the functional form (see supplementary material). We then compared the evidence of the winning models in each analysis; namely, the evidence for the mapping between (sigmoid MRS, self-inhibition), (sigmoid MRS, inter regions), (polynomial MRS, self-inhibition) and (polynomial MRS, inter regions), as shown in Figure 10. The results suggest that the sigmoid transformation is the most likely functional form to explain intersubject variability in the inhibitory recurrent (self) connections in superficial and deep layers.

**Figure 10.**
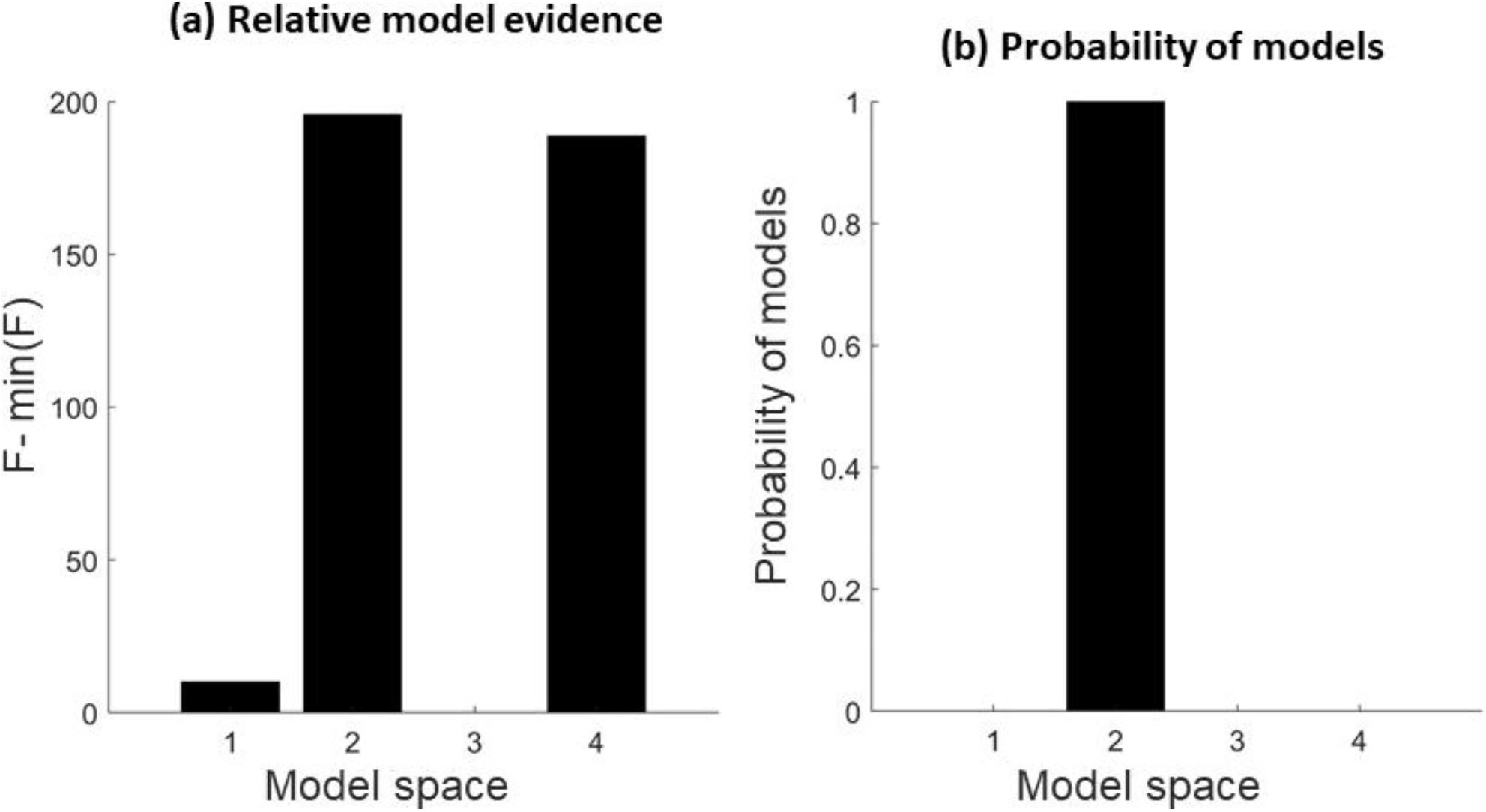
Comparing the probability of sigmoid and second order models that encode the mapping from MRS GABA to self-inhibitory connections (model 2, 4) and interlaminar inhibitory connections (model 1, 3). The comparison indicates the second model is most likely: a sigmoid transformation of MRS GABA informs the recurrent or self-inhibition of deep and superficial populations.

#### 3.2.2 MRS glutamate data as an empirical prior for excitatory synaptic parameters

We performed the proposed analysis to identify the relationship between glutamate concentration and excitatory synaptic connections. We considered different combinations of the excitatory connections over sigmoid and second order polynomial functional forms of the relationship between MRS and synaptic parameters. The correlations of models’ free energy over odd and even data is shown in Figure 11. The winning model over the complete search space confirms that MRS glutamate is linked to excitatory connections from superficial to deep layers and superficial to inhibitory interneurons (model 3).

**Figure 11.**
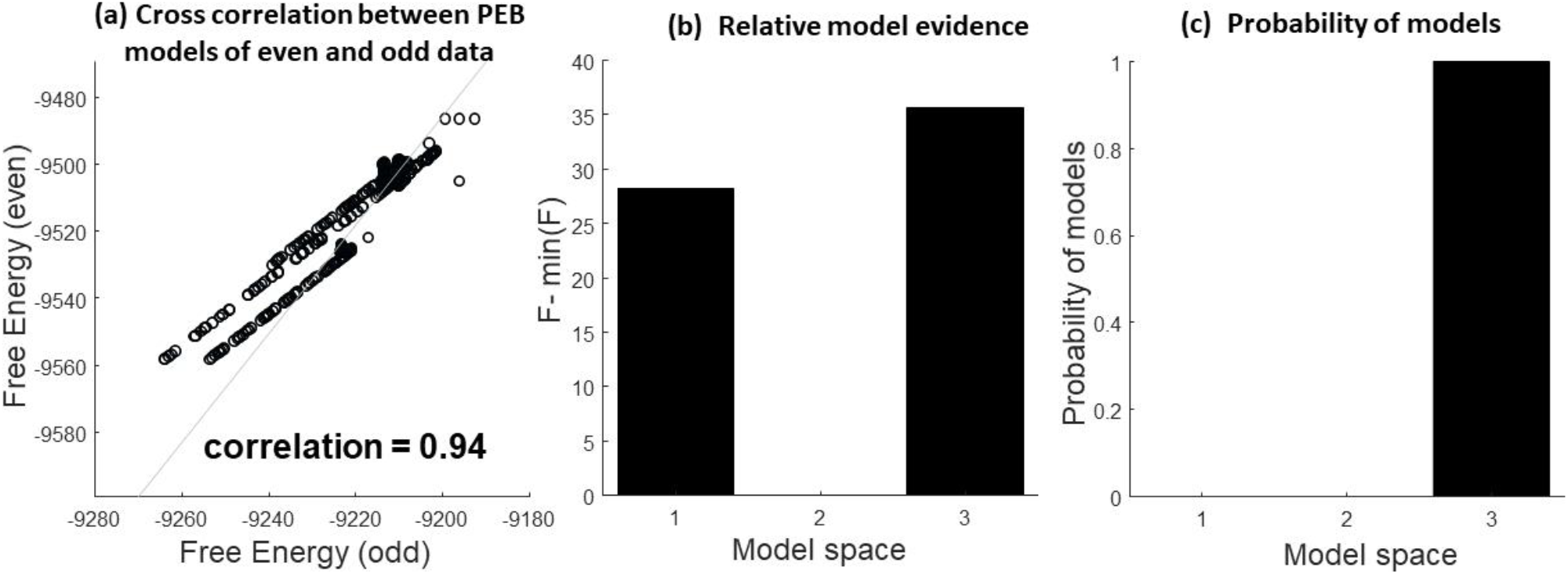
(a) Correlation analysis between the free energy of the PEB model for odd and even data. This plot shows that the PEB approach consistently evaluates model evidence in even and odd data. (b) Free energy and probability of models for the odd and even data are illustrated. Model 1 includes effects on excitatory connections from superficial to deep layers, model 2 considers excitatory connections from superficial excitatory populations to inhibitory interneurons, and model 3 includes both. (c) The transformation of the glutamate that gives the maximum evidence for the odd and even data. Although, there are two MRS functions that maximise model evidence for the odd and even data, both maps suggest similar nonlinear thresholding of glutamate measures provide plausible empirical priors. Please see supplementary Figure 15 for a detailed definition of models in this graphic.

The results of comparing a polynomial form and sigmoid transformation of MRS-glutamate measures suggest that a sigmoid transformation is more likely (see Figure 12 and supplementary material).

**Figure 12.**
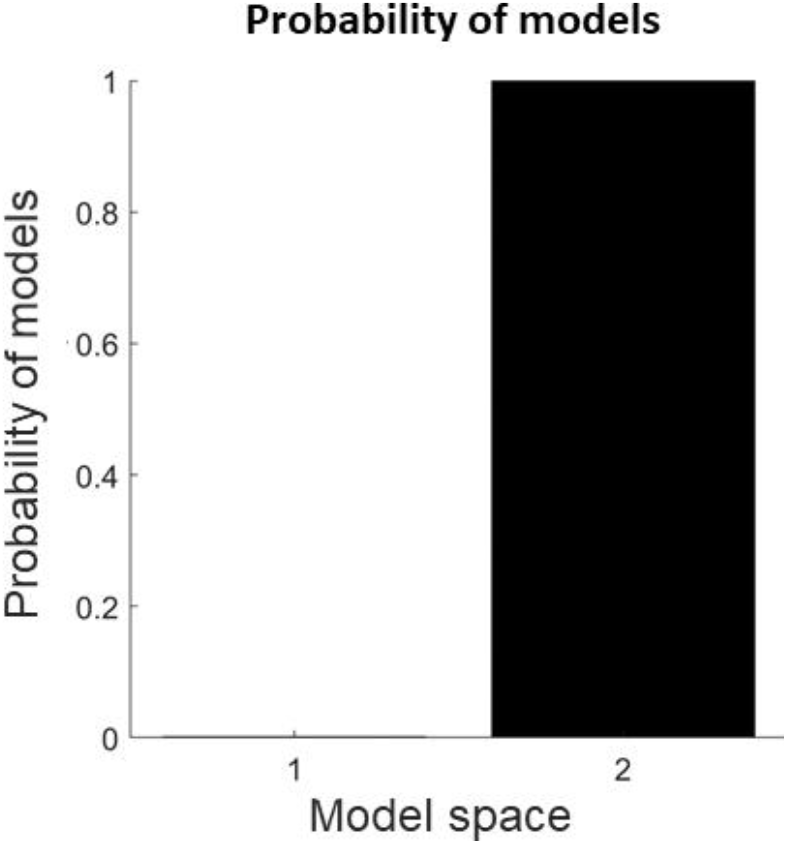
Comparing the probability of models that encode the mapping between MRS glutamate measures and excitatory connections (model 1 is the second order polynomial and model 2 corresponds to the sigmoid function). The comparison indicates model 2 is the most likely: a sigmoid transformation of MRS glutamate measures informs the excitatory connections between deep and superficial layers.

## Discussion

We present a framework for specifying and comparing hypotheses of the relationship between synaptic physiology and neurotransmitter levels, based on a combination of resting-state MEG and MRS. This enables one to specify and compare the effect of biomarkers of inter-subject variation as informative priors on synaptic parameters in a time efficient and Bayes-optimal manner. The method is applicable to generative models of evoked and resting-state time series, recorded by magnetoencephalography, electroencephalography or functional magnetic resonance imaging techniques. In principle, the individual differences specified here by MRS could be replaced by other imaging modalities, or regional neuropathology assays.

We illustrate how the method can be used to test, non-invasively, hypotheses about the relationship between synaptic physiology and neurotransmitter concentrations in humans. This goes beyond the correlation between M/EEG data features and MRS estimates of neurotransmitter concentration (McColgan et al., 2020), in using formal, generative models of the neurophysiological observations. It is computationally efficient, using first level DCM once and re-evaluating the DCM parameters and model evidence analytically using Bayesian model reduction. The model evidence is thereby compared between models with and without the MRS priors, in contrast to the reinversion of the DCMs for alternative priors (Stephan et al., 2009).

We provide evidence that a non-linear transform of the MRS GABA measures offers the best explanation for inter-subject variability in inhibitory recurrent (i.e., self) connectivity of superficial and deep neuronal populations. In addition, a sigmoid transform of the MRS glutamate provides the best explanation of inter-subject variations in excitatory connections from superficial to deep layers. The identification of a sigmoid form for the MRS transform is interesting, because experimental changes of GABA and glutamate concentrations exhibit a sigmoid relationship to neuronal responses, though postsynaptic gains (Benardete and Kriegstein, 2002, Dyke et al., 2017, Chebib et al., 2009). The relationship between superficial and deep layer connectivity is especially relevant to studies of neuropsychiatric and neurological disorders with abnormalities in GABA and glutamate, e.g., schizophrenia, movement disorders and dementia (Adams et al., 2021, Murley et al., 2020, Murley et al., 2022). Using MRS data as an empirical prior on the synaptic connections may provide valuable information about the impact of neuropathology on synaptic function and the response to treatment.

There are several limitations to this study. We focus on the relationship between neurochemical concentrations and synaptic physiology in one cortical region. The choice of the right inferior frontal gyrus was motivated by our interest in frontal lobe function and its impairment in frontotemporal lobar degeneration (Eliasova et al., 2014, Hughes et al., 2015, Murley et al., 2020, Murley et al., 2022) and other neuropsychiatric disorders. However, the method can be applied to multiple regions. For instance, one could combine neurophysiology and spectroscopy from principal nodes in the default mode or salience networks; and test whether MRS data at different regions are associated with extrinsic (between-source) and intrinsic (within-source) connections. Such an approach could test whether local neurotransmitter concentrations in one source influence the activity and connectivity of other sources. The spatial resolution of MRS data is limited. The average neurotransmitter concentrations are captured over multiple cortical columns. Here, the GABA estimate is used as a marker of between-subject differences in neurotransmitters for the lateral frontal cortex. We assume that the voxel-wise estimate approximates the neurochemical concentrations in the neurophysiological source used to extract time series for DCM inversion, noting that the source lies within the MRS voxel (Murley et al., 2020).

Our study size was modest, although similar to (Stephan et al., 2009), where a related method for DCM is described. A larger sample size could widen the intersubject distribution, and facilitate the inversion of the hierarchical models (Kerkhoff and Nussbeck, 2019). However, the Bayes factors indicate that our study had sufficient precision (sufficient ‘power’ by analogy to frequentist testing) to support the inferences made. Although we draw inferences about the role of GABA and glutamate on neurophysiological function, and synaptic connectivity specifically, we do not, in this study, perturb such functions through psychopharmacological challenges. The combination of the current analysis with GABAergic or glutamatergic interventions could be used to identify baseline dependent effects of drug interventions, as in (Adams et al., 2021) within a simpler and integrated Bayesian modelling procedure.

In conclusion, we propose that dynamic causal models of neurophysiology can be explicitly informed by priors based on measures of individual differences in neurochemistry, molecular pathology or cell/synapse specific loss. The enrichment of DCMs by such markers of inter-subject variability has many potential applications, exploiting the computational efficiencies of parametric empirical Bayesian methods and Bayesian Model Reduction with hierarchical inversion of individual and group-level models of functional imaging data.

## Acknowledgements

This work has been funded by the Wellcome Trust (220258), the Medical Research Council (SUAG/092 G116768) and Cambridge Centre for Parkinson-plus; the Holt fellowship; and the NIHR Cambridge Biomedical Research Centre (BRC-1215-20014). KJF is supported by funding for the Wellcome Centre for Human Neuroimaging (Ref: 205103/Z/16/Z) and a Canada-UK Artificial Intelligence Initiative (Ref: ES/T01279X/1). The views expressed are those of the author(s) and not necessarily those of the NIHR or the Department of Health and Social Care. For the purpose of open access, the authors have applied a CC BY public copyright licence to any Author Accepted Manuscript version arising from this submission.

## 4 Supplementary figures

Figure 13 shows the first level DCM results across all eleven participant. Figure 14 illustrates model comparison of second level PEB associated with self-inhibitory connections (top) and inter-regional connections (bottom plot) given the class of first and second order polynomial maps for MRS data as their regressors. Figure 15 illustrates the definition of models in Figures 7, 9 and 11.

**Figure 13.**
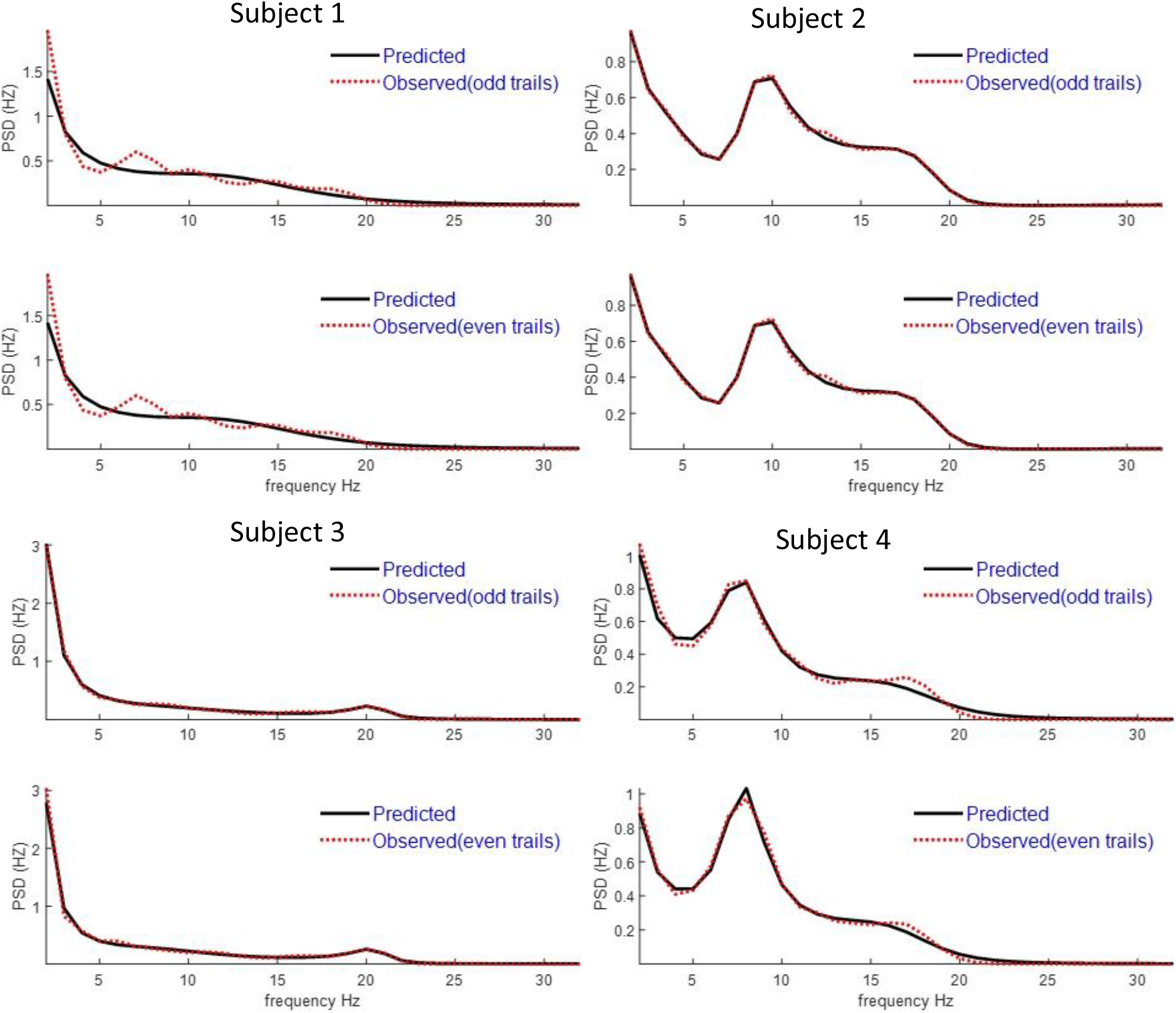
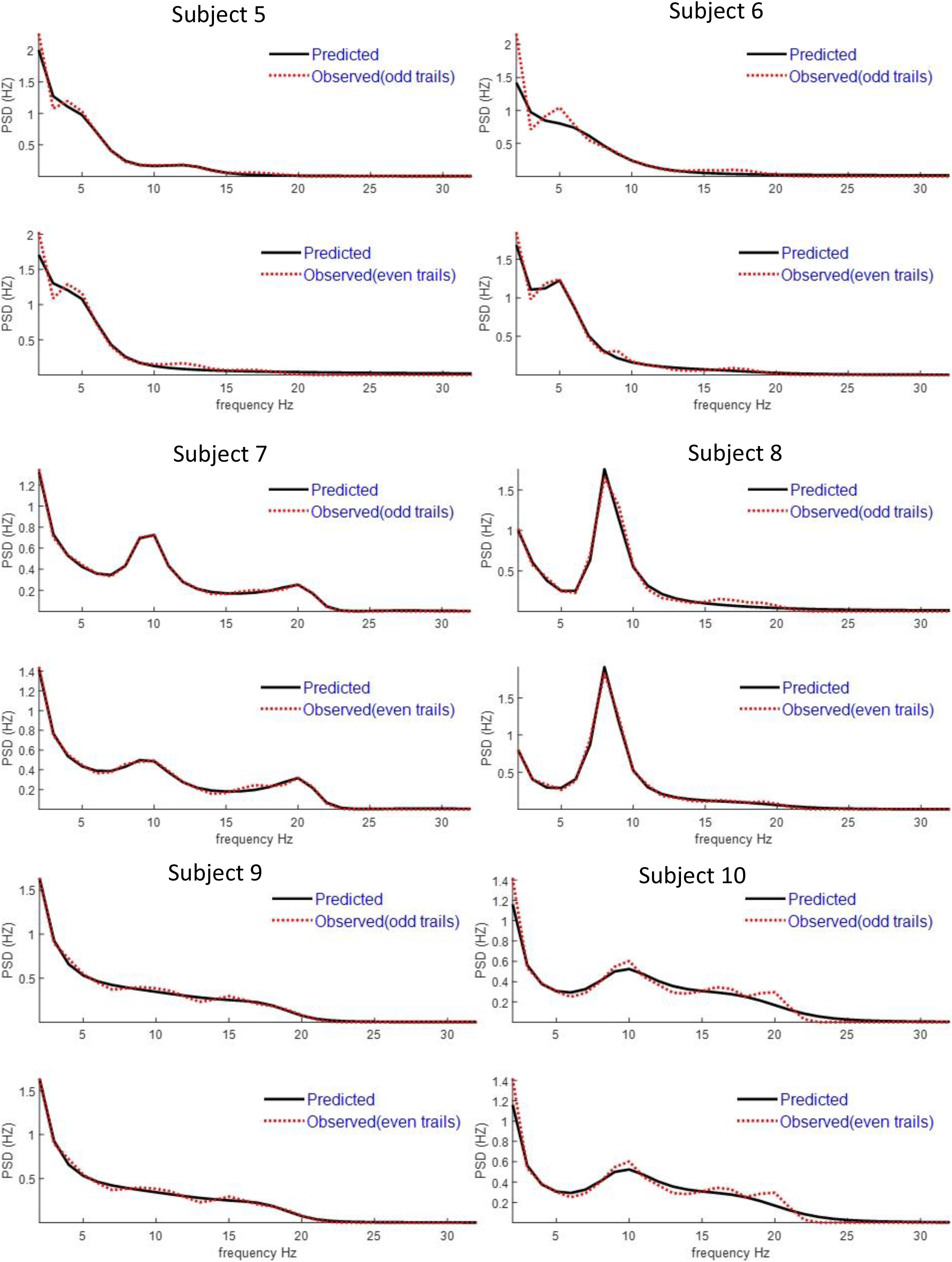
First level DCM predicted vs observed data for each subject even and odd PSD data. The variance explained across all data (even/odd) is 98%.

**Figure 14.**
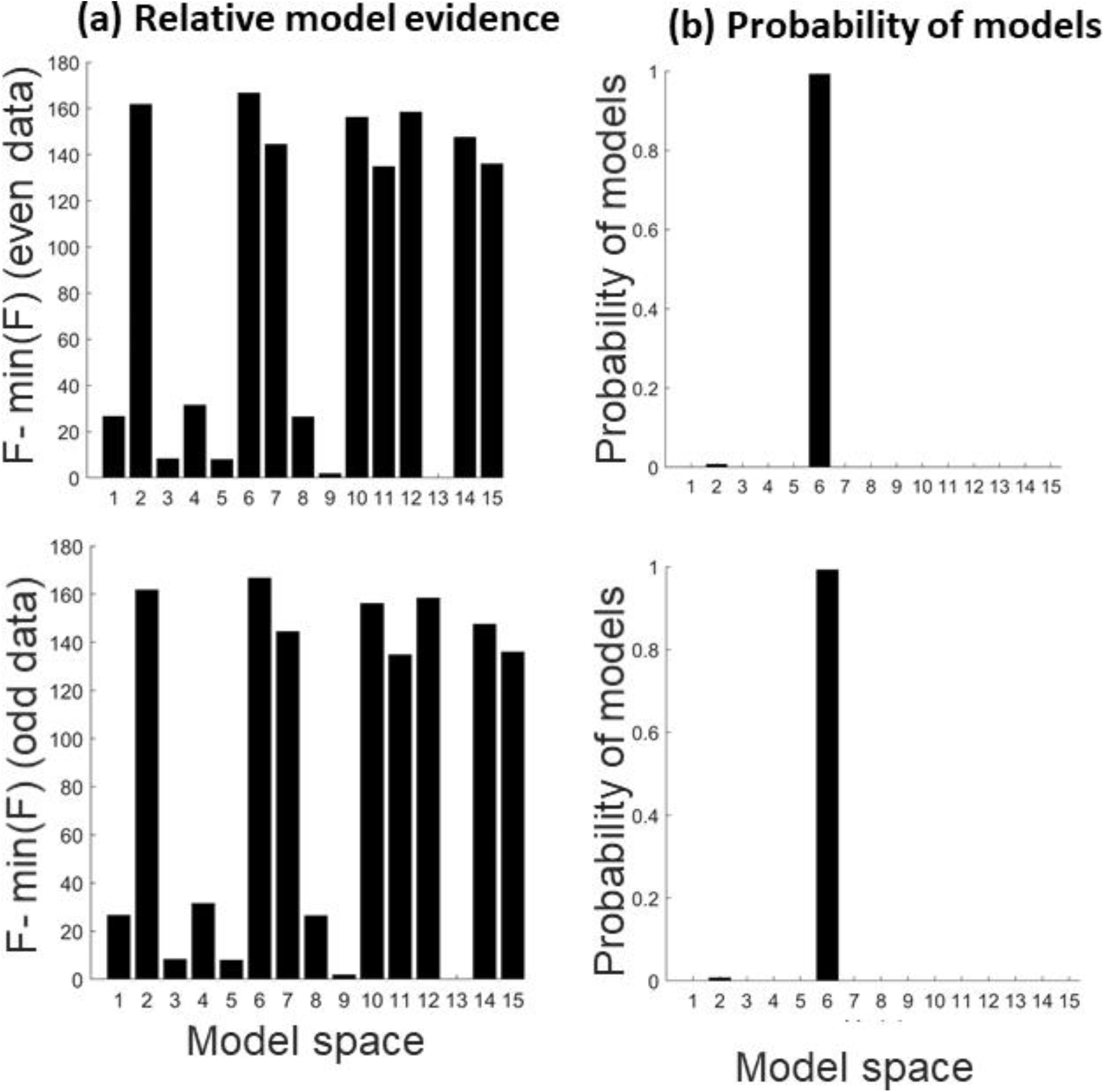
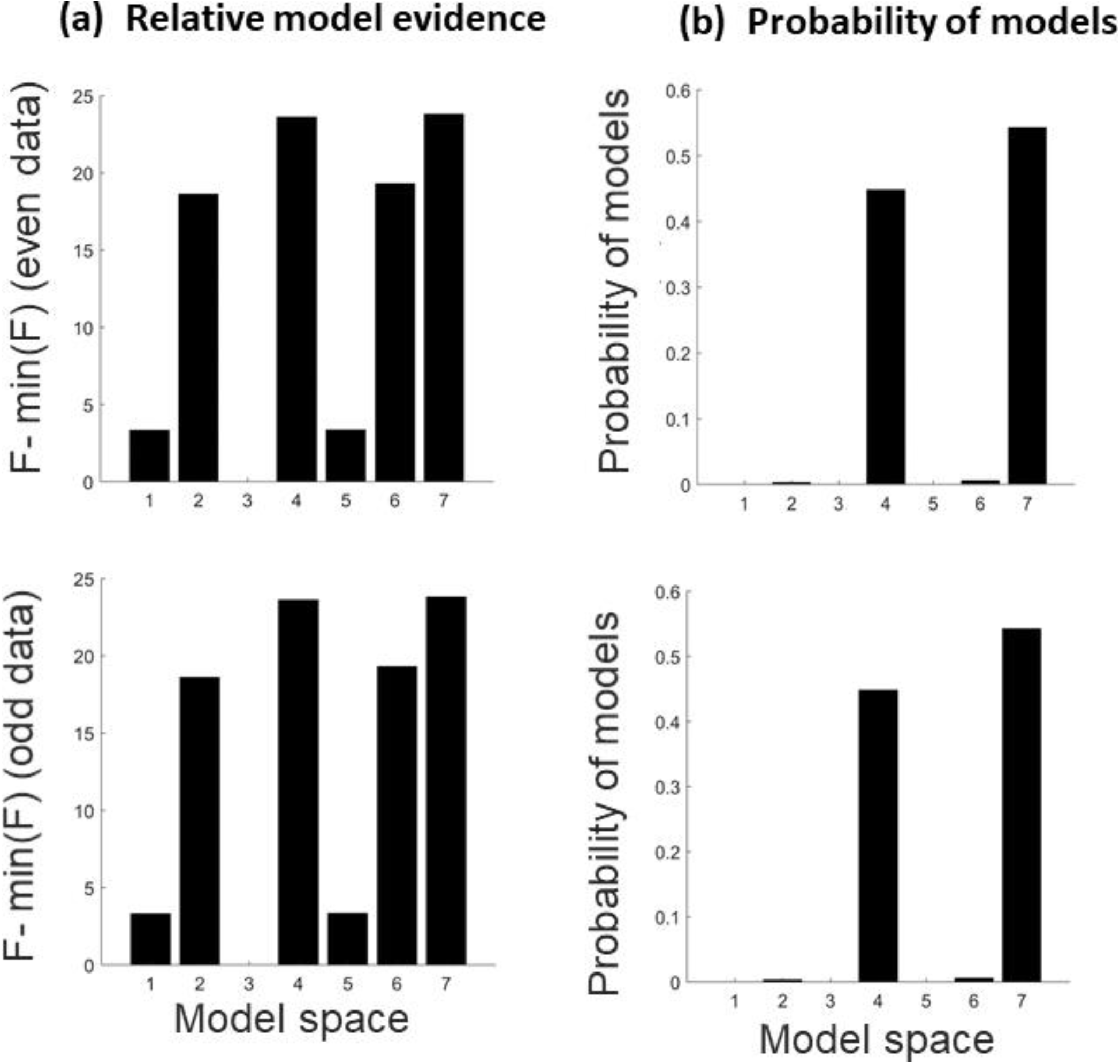
The following figure shows the second level PEB model comparison results with MRS maps (class of first and second order polynomial) as their regressors. The model comparison associated with different combinations of self-inhibitory connections are in the top plot and the model comparison associated with different combinations of inter-regional connections are in the bottom plot.

**Figure 15.**
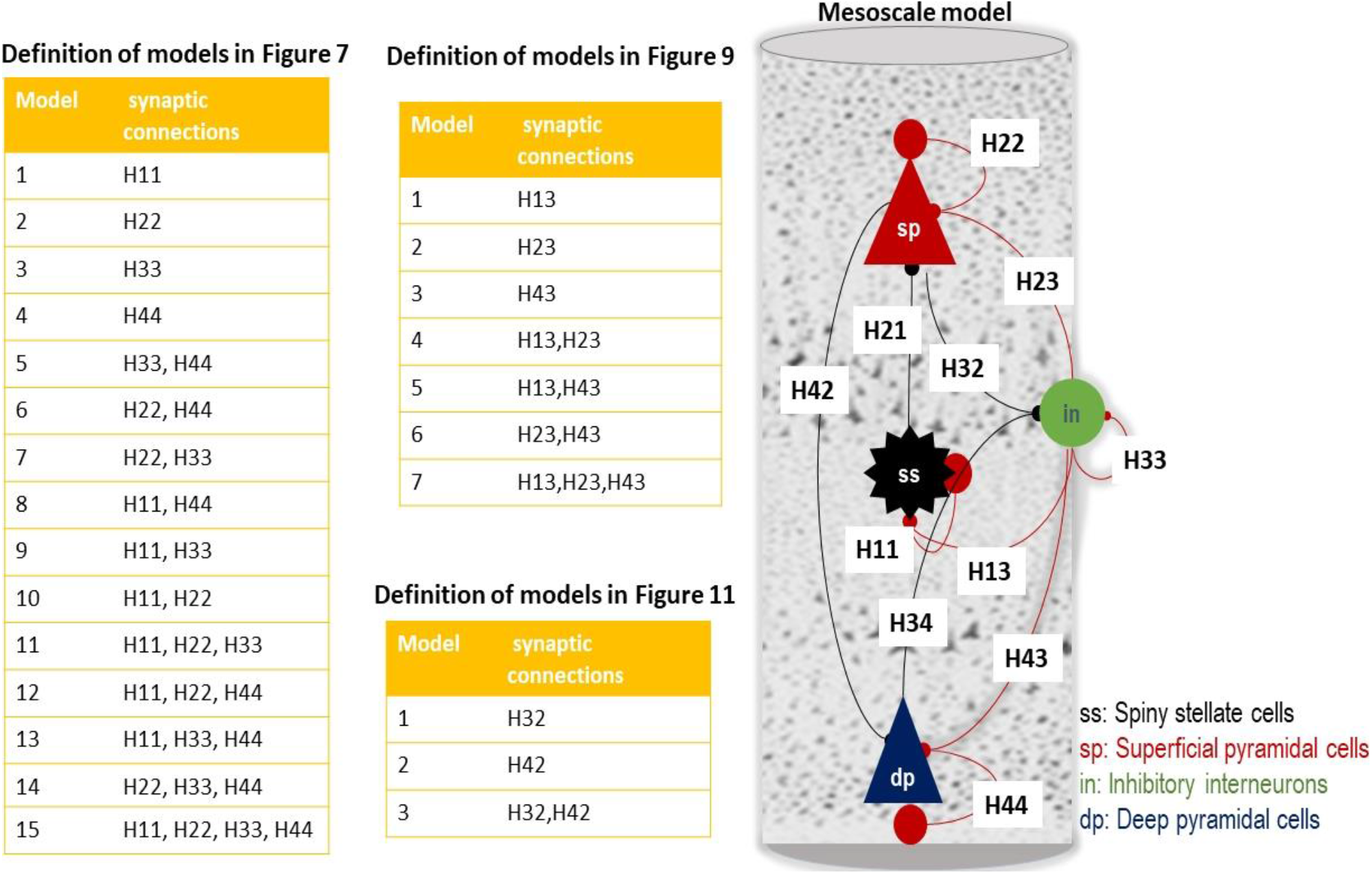
Definition of models in Figures 7, 9 and 11. This graphic illustrates which synaptic connections are informed by MRS data in Figures 7, 9 and 11. The right hand-side graphics show different synaptic connections in the conductance based models.

